# Elucidating multi-input processing 3-node gene regulatory network topologies capable of generating striped gene expression patterns

**DOI:** 10.1101/2021.11.01.466847

**Authors:** Juan Camilo Arboleda-Rivera, Gloria Machado-Rodríguez, Boris A. Rodríguez, Jayson Gutiérrez

## Abstract

**Background:** A central problem in developmental and synthetic biology is understanding the mechanisms by which cells in a tissue or a Petri dish process external cues and transform such information into a coherent response, e.g., a terminal differentiation state. It was long believed that this type of positional information could be entirely attributed to a gradient of concentration of a specific signaling molecule (i.e., a morphogen). However, advances in experimental methodologies and computer modeling have demonstrated the crucial role of the dynamics of a cell’s gene regulatory network (GRN) in decoding the information carried by the morphogen, which is eventually translated into a spatial pattern. This morphogen interpretation mechanism has gained much attention in systems biology as a tractable system to investigate the emergent properties of complex genotype-phenotype maps.

**Methods:** In this study, we apply a Markov chain Monte Carlo (MCMC)-like algorithm to probe the design space of three-node GRNs with the ability to generate a band-like expression pattern (target phenotype) in the middle of an arrangement of 30 cells, which resemble a simple (1-D) morphogenetic field in a developing embryo. Unlike most modeling studies published so far, here we explore the space of GRN topologies with nodes having the potential to perceive the same input signal differently. This allows for a lot more flexibility during the search space process, and thus enables us to identify a larger set of potentially interesting and realizable morphogen interpretation mechanisms.

**Results:** Out of 2061 GRNs selected using the search space algorithm, we found 714 classes of network topologies that could correctly interpret the morphogen. Notably, the main network motif that generated the target phenotype in response to the input signal was the type 3 Incoherent Feed-Forward Loop (I3-FFL), which agrees with previous theoretical expectations and experimental observations. Particularly, compared to a previously reported pattern forming GRN topologies, we have uncovered a great variety of novel network designs, some of which might be worth inquiring through synthetic biology methodologies to test for the ability of network design with minimal regulatory complexity to interpret a developmental cue robustly.

**Author summary:** Systems biology is a fast growing field largely powered by advances in high-performance computing and sophisticated mathematical modeling of biological systems. Based on these advances, we are now in a position to mechanistically understand and accurately predict the behavior of complex biological processes, including cell differentiation and spatial pattern formation during embryogenesis. In this article, we use an *in silico* approach to probe the design space of multi-input, three-node Gene Regulatory Networks (GRNs) capable of generating a striped gene expression pattern in the context of a simplified 1-D morphogenetic field.

## Introduction

Cells interact continuously with their environment, which requires precise regulatory strategies to avoid potential detrimental responses. Arguably, most cellular functions arise from the dynamic activity of Gene Regulatory Networks (GRNs), which play a central role in interpreting external and internal signals. This information processing function is critical in developmental processes such as those in which a group of cells differentiates in response to a signaling molecule. Such molecules were referred to as morphogens by Turing in 1952 [1], and posterior theoretical studies on patterning led to the conceptualization of the French Flag Problem by Wolpert, who also stated in this respect that a gradient of concentration of a morphogen could trigger cell differentiation in a one-dimensional field of cells [2, 3]. Although the Bicoid protein (bcd), the first example of a molecule that acted as a morphogen, was only found in the 80s in the developing embryos of *Drosophila melanogaster* [4, 5], now it is known that there are many other examples such as the Decapentaplegic (Dpp) protein in *Drosophila* wing imaginal discs [6], as well as Sonic Hedgehog [7, 8] and Wnt [9]. However, information processing by GRNs has proven to be a complex process, and a major goal of developmental biology is to understand mechanistically how positional information conveyed by morphogens is translated into spatial differentiation.

Our understanding of how GRNs process information has increased thanks to the concept and theory of network motifs, defined as patterns of interconnections occurring in networks at numbers significantly higher than those in randomized networks [10]. This novel approach, along with mathematical modeling and computational systems biology methods, is a powerful tool to study GRNs [11], and experimental studies have validated the predictions of studies using these methods [12–15]. Some have even used synthetic GRNs to produce artificial cell differentiation [16], circadian gene expression [17], counting devices [18] and systems that respond to light [19, 20]. Moreover, the design and implementation of GRNs through synthetic biology is emerging as a promising tool to study biological phenomena as pattern formation [21] and as a novel therapeutic tool with interesting biomedical applications [22–26].

In a tissue context, the generation of a stripe of gene expression is a fundamental patterning function in development, and it has been shown that simple feed-forward motifs can robustly achieve such a patterning task [27]. For example, Cotterell and Sharpe studied what kind of 3-node network topologies could effectively translate a morphogen gradient into a striped gene expression pattern in a one-dimensional field of cells. They found a variety of networks that implemented at least 7 different mechanisms with varying complexity levels, most of them variations of feed-forward motifs [28].

Although GRNs with positional information processing capacities have been extensively analyzed, most studies have so far emphasized particular regulatory systems that respond to a single input only. These studies have typically focused on either regulatory systems observed in nature [29–31], synthetic implementations with predefined topologies [16, 32–34], or have deliberately constrained the study of GRNs to just a handful of alternative designs [27, 35]. However, previous work has demonstrated the necessity to expand computational analysis of pattern forming GRNs to multi-input settings. For instance, it has been shown that the neural subtypes specification system in the vertebrate neural tube involves a two-input network in which Sonic Hedgehog acts as a morphogen and the Olig2 and Nkx2.2 genes can act as the receiver nodes [7, 36, 37].

In this study, we apply a Markov chain Monte Carlo (MCMC)-like algorithm to probe the design space of three-node GRNs looking for topologies capable of translating a morphogen gradient (input signal) into a striped pattern of gene expression (phenotype). Importantly, unlike most previous computational studies, here we allowed any node (e.g., gene) in a GRN to perceive the input signal in varying ways and selected the best-performing GRNs based on a fitness criterion used to assess the quality of the phenotype with respect to a prescribed optimal pattern. Based on this computational strategy, we uncovered a great variety of distinct classes of network topologies that tend to form a complex interconnected meta-graph that could be easily traversed throughout evolution via single changes in the wiring of the different GRNs.

## Materials and methods

### Gene Regulatory Networks and morphogenetic field

In order to study three-node GRNs we represented them by three sets of real numbers, the first set was composed of the interaction values between genes in the network; the second of the diffusion rates (D) of each of the three gene products; and the third of the degradation rates (*δ*) of these same gene products, with *D* ∈ [0, 0.1] and *δ* = log_2_ *phl*, where *phl* ∈ [5, 50] is the gene product half-life.

The set of interaction values was represented as adjacency matrices *W* ∈ ℝ^3×4^ where *i* represents a regulated node of the GRN and *j* represents a regulator node of the GRN (Fig 1A and 1B) with *w_ij_* ∈ [−10, 10] (this interval was arbitrarily chosen and it is inherited from the model of Munteanu et al. [27] and Cotterell & Sharpe [28]). Values in these matrices can be negative, zero or positive and represent represion, no interaction, or activation respectively. The magnitude of the value is proportional to the strength of the interaction; however, this and all other magnitudes in our model are dimensionless. In this study we define a ‘genotype’ as the set of interaction, diffusion and degradation values that represent a GRN.

**Fig 1.**
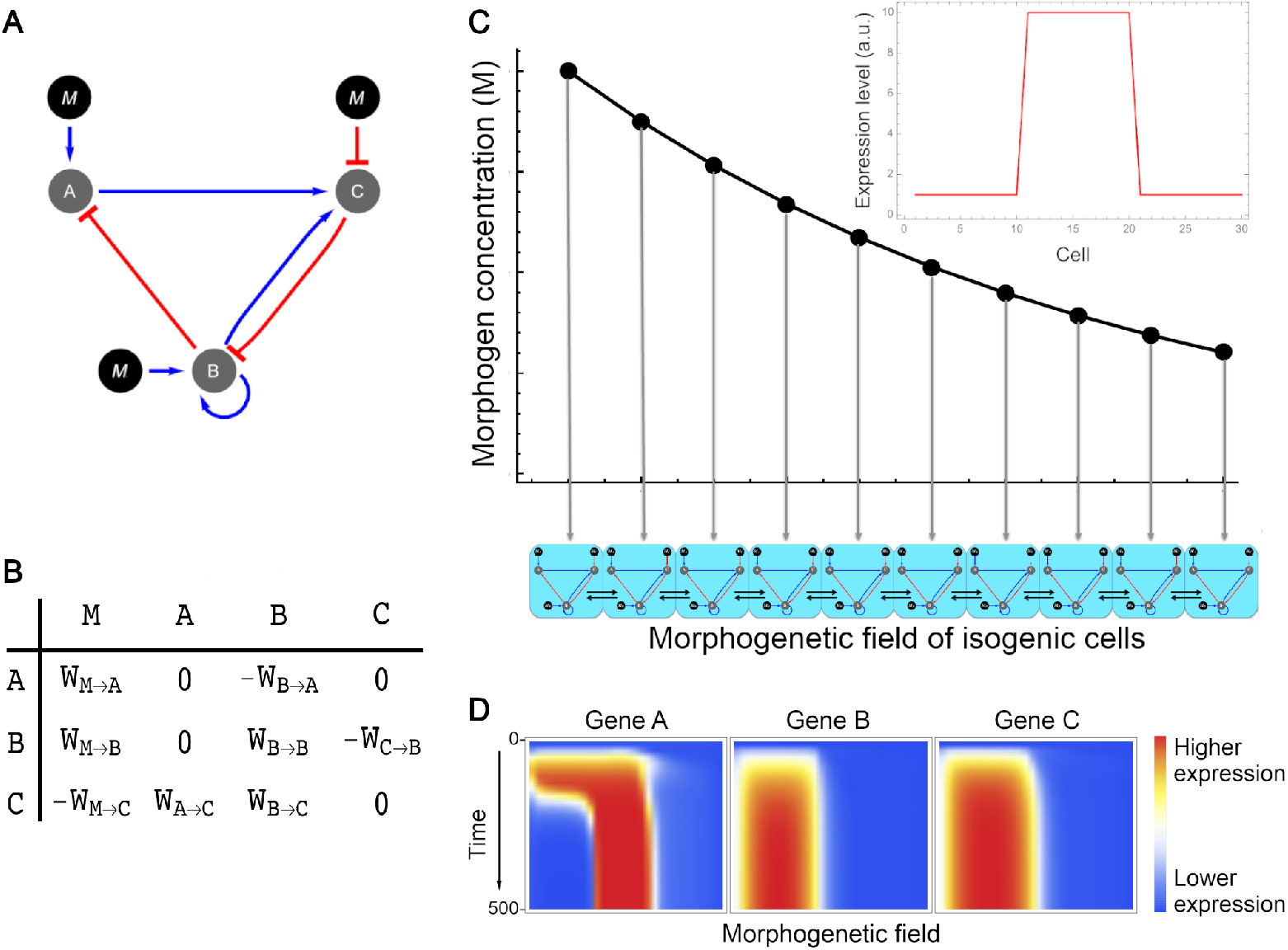
Modelling of GRNs. (A) GRN represented as a directed graph with activation (blue arrows) and inhibition (red arrows) interactions. Black circles tagged with a **M** represent the morphogen; **A**, **B** and **C** represent genes in the network. (B) The same GRN represented as an adjacency matrix in which positive values represent activation, negative ones represent inhibition and zeros represent no interaction. (C) Morphogenetic field. In our model the field was composed of a linear array of 30 isogenic cells and a morphogen gradient concentration described by an exponential decay function. Horizontal arrows between cells represent diffusion of gene products between adjacent cells. Optimal gene expression pattern is shown top right. (D) Heatmap of spatiotemporal expression profile of a GRN where blue colors represent lower expression and red colors represent higher expression. Time axis goes from 0 to 500 integration steps. Horizontal axis represents the morphogenetic field, i. e. cells from 1 to 30 for each gene.

In our model the morphogen could interact with any of the three genes on the GRN, but could not be affected by them. The morphogenetic field was defined by an unidimensional array of 30 isogenic cells exposed to the morphogen concentration gradient (Fig 1C). In multiple-input networks, all the genes of a given cell are exposed to the same morphogen concentration, given by an exponential decay function (see Morphogen spatial distribution). The initial concentration of each gene product in all cells was set to 0.1 in all simulations.

### Mathematical model

Our model is a modification of the model used by Cotterell & Sharpe [28] and proposed by Reinitz et al. [30]. The model is a dynamical system that describes the change in concentration of gene product *i* in time, as shown in Eq 1.

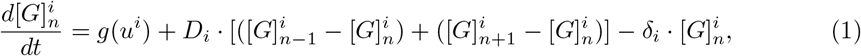

in which 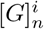 is the concentration of the i-th gene in the *n*-th cell 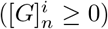, *g*(*u^i^*) is a function describing the relationship between the interactions on the *i*-th gene and its expression Eq (2) and described in more detail below. *D_i_* and *δ_i_* are the diffusion and the degradation rate of *i*-th gene product, respectively.

Sigmoid functions are often used to approximate Hill functions describing gene activation/inhibition in which steepness can be modulated by mechanisms as molecular cooperativity and target sequestration [38]. This approach has been used extensively to model the response of signaling pathways and gene interactions in GRNs [39–42]. Moreover, differential equation models using sigmoid functions have shown to fit gene expression data [43, 44]. In this study, the input function representing regulatory interactions is a sigmoid function described by Eq 2.

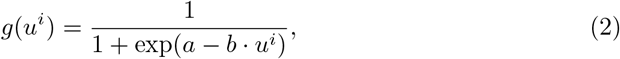

in which *a* is the sigmoid steepness, equal to 5; *a/b* is the threshold value, set to 1 for all simulations, and *u^i^* is the following equation:

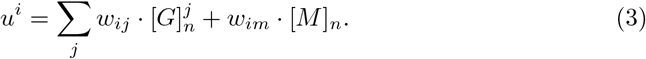

This equation sums the interactions acting upon the *i*-th gene, being *w_ij_* the interaction strength of the *j*-th gene upon the *i*-th gene and *w_im_* the interaction strength of the morphogen upon the *i*-th gene. 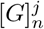 and [*M*]_*n*_ are the concentrations of the *j*-th gene product and the morphogen in the *n*-th cell.

### Morphogen spatial distribution

The morphogen concentration along the morphogenetic field is described by Eq 4.

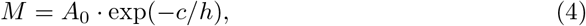

were *A*_0_ is the concentration of the morphogen in the position zero of the morphogenetic field and was set to 1 in our experiments; *c* is the cell index, defined as the ratio between the *n*-th cell from left to right and the total number of cells, and *h* is a decay parameter, whose value in our model was set to 0.4. We do not consider here how the morphogen gradient can be formed, i.e., we do not consider morphogen dynamics but we assume a static gradient.

The phenotype of a GRN was defined as the expression pattern of each gene along the morphogenetic field after 500 time steps of integration of the dynamical system. We chose that number of time steps because numerical experiments showed that GRNs reached the steady state in approximately 300 steps (S1 Fig).

### Optimal pattern definition

The optimal pattern of gene expression defined in this study consists in cells at the border of the field (*n* < 11 and *n* > 20) displaying expression levels lower than 10% of the maximal level observed along the field of cells for the output gene, and cells at the middle of the field (*n* ∈ [11, 20]) displaying expression levels greater than 90% of the maximal level observed along the entire field for the output gene (Fig 1C).

### Search space algorithm

The Markov chain Monte Carlo (MCMC)-like algorithm used to produce the set of gene regulatory networks (GRNs) consisted in the following steps:

1. **Generate a random GRN.**
2. **Numerically solve the dynamical system for the GRN from t=0 until t=500.** We integrated the differential equation using the function NDSolve from Wolfram Mathematica 11 with the option “EquationSimplification” and the “Residual” simplification method.
3. **Evaluate fitness by comparing the GRN phenotype with the optimal phenotype (Fig 1C).** In order to compute the fitness function for each GRN we used three different filters. The first filter assesses whether the expression profile of the output node reaches a quasi-steady state:

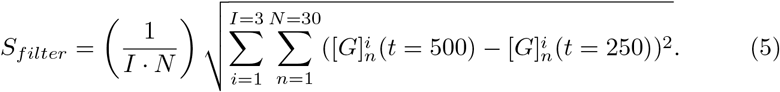 The expression profile was considered to have reached the steady state if *S_filter_* < 0.001. The second filter measures the spatial heterogeneity in the field:

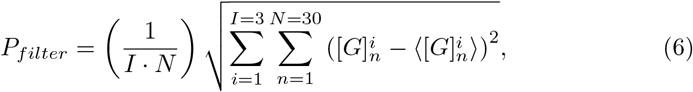

where 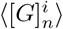 is the mean concentration of the *i*-th gene along the field at *t* = 500. The third filter relates the Manhattan Distance (*D_obs_*) between an expression pattern and the optimal pattern with the maximum distance achievable (*D_max_*):

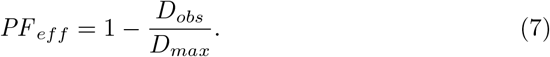 In this function, the expression profile of the output gene was normalized and discretized so that each expression value for each cell in the field was an integer ranging from 1 to 10. The filters mentioned above are integrated in the following functions that evaluate the quality of a given phenotype:

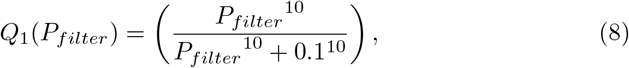

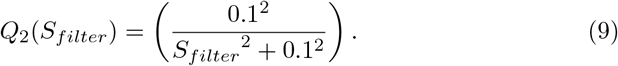 Finally, these quality functions were used to compute the fitness score with the following fitness function:

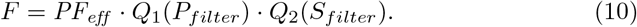
4. **Mutate GRNs randomly and evaluate fitness again.** Three types of mutations were possible in our algorithm: 1) changes in the numerical value of the degradation parameter, 2) changes in the numerical value of the diffusion parameter and 3) changes in the numerical values of the interaction strengths. The type of mutation to be performed was selected randomly with a probability of 0.15 for type 1 and type 2 mutations and 0.7 for type 3 mutations. If type 3 mutation was selected, either the interaction strength was set to zero (with a probability of 0.2) or was changed randomly (with a probability of 0.8). If fitness of the mutated GRN was greater or equal than that of the previous GRN, the mutated GRN was saved and the cycle continued from step 2. This cycle was repeated 50000 times, and a GRN sampled at a given step and with a fitness score ≥ 0.95 was saved to a file as long as its associated topology had not been sampled in previous steps.

Steps 1 to 5 were iterated 500 times, resulting in a set of 2061 GRNs with a striped pattern of gene expression in at least one gene.

The source code for the search space algorithm and other parts of analysis is available at https://github.com/Nesper94/gene-regulatory-networks.

### Classification of GRNs

In order to group all the isomorphic (i.e. topologically equivalent) networks from the totality (2061) of sampled GRNs we used a classification algorithm that selected a GRN and generated all isomorphs of that GRN by interchanging rows of the adjacency matrix (*i* ↔ *j*) while interchanging the corresponding columns (*i* + 1 ↔ *j* + 1). This kind of transformation of the adjacency matrix results in isospectral graphs that are also isomorphic to the original graph and is equivalent to generate a permutation matrix *P* and multiplying *PMP^T^*, where *M* is the adjacency matrix and *P^T^* the transpose of *P*. The algorithm then compared these isomorphs with another GRN and classified the topologies in the same group if at least one of the transformed adjacency matrices (or the original one) was identical to that of the other GRN, i. e., if they were isomorphic networks.

### Neutral network

The neutral network was obtained by generating all possible isomorphs of a network topology and then calculating the minimal Hamming distance between these isomorphs and other network topologies. The Hamming distance between two topologies was calculated using the following equation:

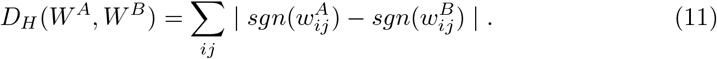

We adopted the same definition of neighborhood as Cotterell & Sharpe [28], in which two topologies were neighbors if the gain or removal of any one interaction can transform one of the topologies into the other.

### Robustness analysis

In order to evaluate the robustness of the different topologies to perturbations in parameters of interactions between genes we took a GRN belonging to each one of the network topologies and we modified each interaction independently by increasing and decreasing its value by 20%. For topologies with 3 or more GRNs we chose the most similar to the mean configuration for that topology.

As a measure of the robustness of the topology we chose the proportion of times in which a perturbation resulted in a GRN with a fitness greater or equal to 0.95. Additionally, we considered a topology to be robust if the previous measure was greater or equal to 0.5.

As a measure of the robustness of subgraphs (see “Classification of topologies”) we calculated the mean of the robustness measure of all the topologies containing a particular subgraph.

### Robustness of topology 9 to perturbations in non-network parameters

In order to estimate the robustness of topology 9 to changes in parameters that control morphogen gradient and changes in initial concentrations of gene products, we generated 100 fitness values for each one of the 19 GRNs belonging to topology 9; in each of these 100 assays we chose *A*_0_, *h* or initial concentrations randomly from a Normal Distribution with *μ* = 1 for *A*_0_, *μ* = 0.4 for *h* and *μ* = 0.1 for initial concentrations and a Coefficient of Variation of 30% in each case.

### Shannon entropy

We calculated the Shannon entropy Eq (12) of the network topologies abundance distribution in order to test if this distribution could be obtained by chance with a probability greater than 0.05. In order to obtain a 95% confidence interval, we generated 30 samples of a set of 2061 random matrices and calculated the Shannon entropy in bits for each sample and tested these data for normality.

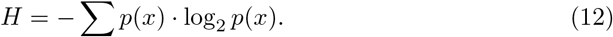

### Classification of GRNs phenotypes

In order to classify each GRN by its spatiotemporal expression profile (its dynamics of expression), we selected expression profiles of GRNs from time step 0 to time step 250 each 10 time steps, for a total of 26 expression profiles over time. These spatiotemporal expression profiles were represented as tensors *S* ∈ ℝ^3×26×30^, in which each element (*s_itn_* ≥ 0) is the expression level of the *i*-th gene at time *t* in cell *n*.

Next, we calculated the distance between all the pairs of tensors and performed a Neighbor Joining clustering using the package Scikit Bio 0.5.5 from Python 3 [45].

### Classification of topologies

In order to classify each topology by its subgraph composition, we first created a matrix *T* = (*t_ij_*) with each row corresponding to a different topology and 17 columns corresponding to subgraphs shown in S2 Fig. Each element *t_ij_* of the matrix was 1 if the *j*-th subgraph was present in the *i*-th topology or 0 if the subgraph was not present.

With this matrix we then calculated a distance matrix in which each element consisted in the Hamming distance between the row vectors in matrix *T* between each pair of topologies. We then clustered the topologies based on the Neighbor Joining method, as implemented in the Scikit Bio package [45].

### Complexity index and network descriptors

As a measure of network complexity, we used the complexity index based on Shannon entropy proposed by Bonchev & Rovray [46]. We calculated this index using the following equation:

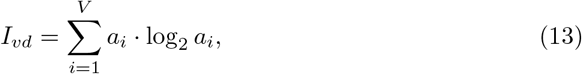

where *a_i_* denotes the degree of the *i*-th node and *V* the number of nodes in the network. Both the network descriptors and Pearson’s chi-square goodness of fit tests were performed for the node degree distribution using the built-in functions from the Wolfram Mathematica software.

### Pattern formation under varying size of the morphogenetic field

We evaluated the ability of each of the sampled GRNs to generate a striped pattern of gene expression under varying sizes of the morphogenetic field. Specifically, we calculated the fitness of each GRN across morphogenetic fields composed of 10, 20, 40, and 50 cells. In these simulations we redefined the morphogen gradient in such a way that the initial and final concentration were constant in all the fields (S3 FigA). In order to know if GRNs with the same topology had a similar fitness in the different fields, we performed a Principal Component Analysis in Scikit-learn 0.24.2 [47] using the fitness values of each in the four different fields (see S3 Fig).

## Results

The classification of GRNs showed that our initial set of 2061 GRNs can be grouped into 714 distinct network topology classes, with each containing a variable number of GRNs (Fig 2). This abundances distribution was not produced by chance as its Shannon entropy was significantly lower than that of random networks produced without selection involved (Shannon entropy = 8.52963, 95% confidence interval for random networks = [10.8915, 10.9275]), and instead it displays some level of structure, indicating the existence of network motifs.

**Fig 2.**
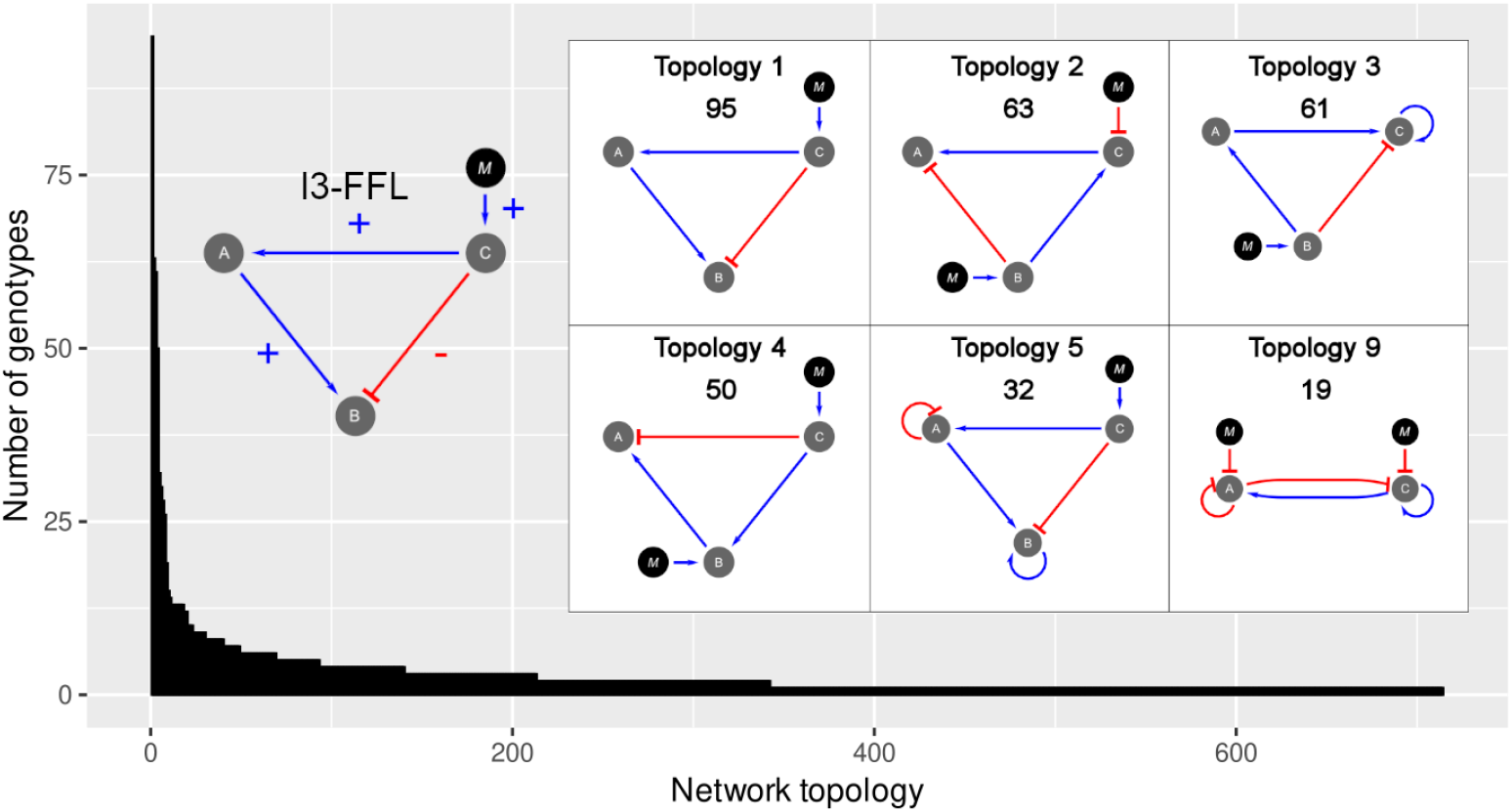
Distribution of abundances of network topologies. Some of the most frequent network topologies shown in the grid contain the I3-FFL network motif (shown on the left). In this network motif the gene B is activated in an indirect way by the morphogen through the activation of A (positive interactions), and at the same time, it is inactivated in a direct way by the morphogen (negative interaction). Topologies are numbered in order of decreasing abundance, being the Topology 1 the most abundant with a number of 95 genotypes. The number of genotypes belonging to each network topology is presented below the topology number. Note that network topology number 9 produces the striped phenotype using a two-node GRN.

The most abundant network topologies are shown in Fig 2. Interestingly, among the 15 most abundant network topologies, eleven of them presented the Incoherent type 3 Feed-Forward Loop (I3-FFL) network motif. In addition, we found that topology number 9 could produce the striped phenotype with only two nodes.

The neutral network (a metagraph in which each node is a network topology) shows that most of the network topologies are grouped into a connected graph. This graph is composed of 639 nodes that are accessible by one mutational step. The 15 most abundant network topologies, with the exception of number 10, are part of this graph (Fig 3). We also found nodes inside the neutral network that had more connections between them than with other nodes, which leads to further clustering of nodes. Interestingly, in almost every cluster one can find one or more of the 15 most abundant network topologies. In addition, we found that the first eight most abundant network topologies were located in the largest cluster.

**Fig 3.**
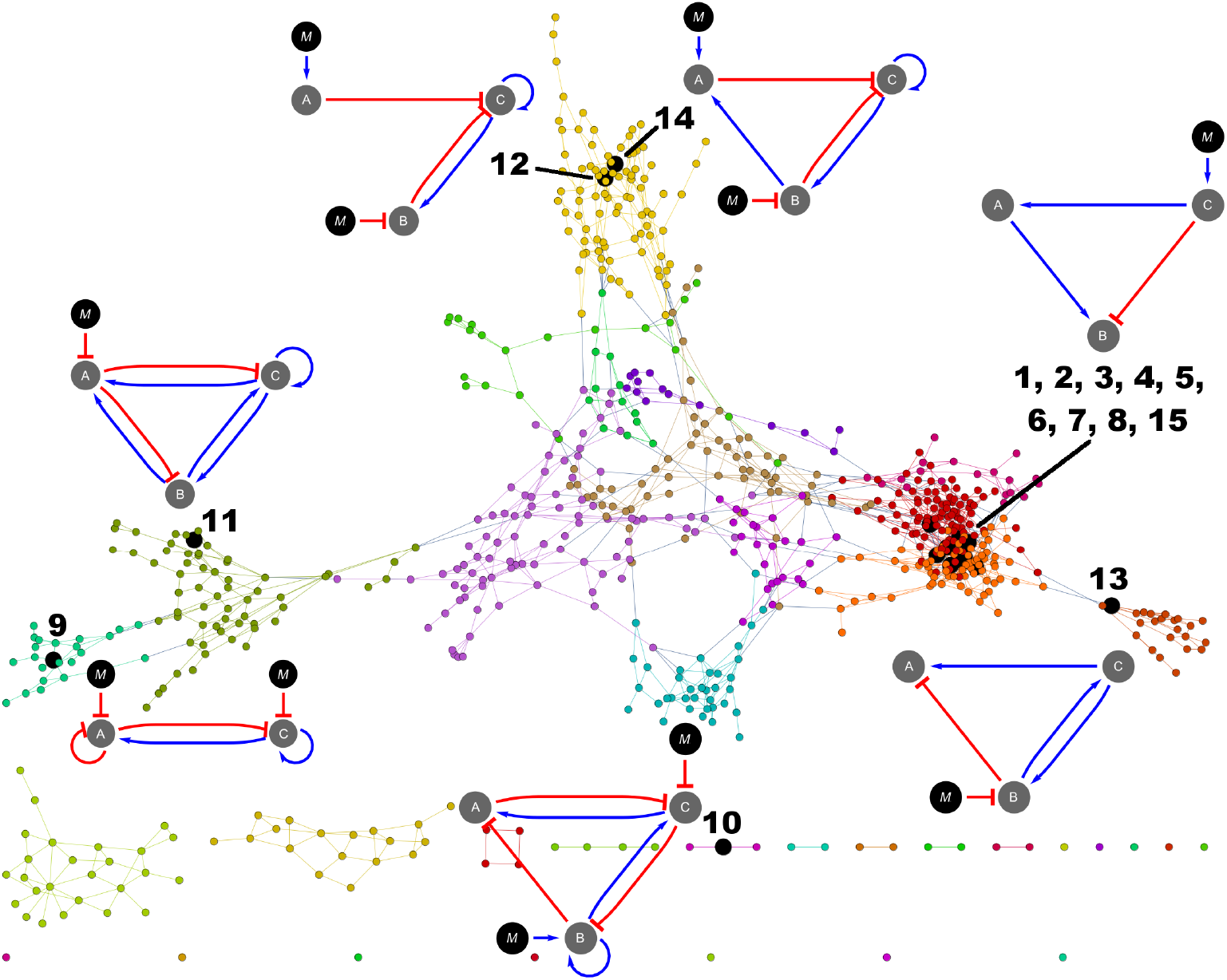
Neutral network. This graph shows network topologies connected by one mutational step, where each node represents a single topology. Black nodes represent the most abundant topologies. All nodes with the same color belong to a cluster of nodes more connected between them than with other nodes in the graph. The most abundant topologies are represented as directed graphs in the same way as in Fig 1A, except for the topologies 1 to 8 and 15 (orange and red clusters), in which only topology 1 is depicted.

The degree distribution of this neutral network seems to follow a Poisson distribution (Pearson’s *χ*^2^ = 12.8902, p-value = 0.115684) characteristic of a Erdös-Rényi network with many nodes and edges [48]. This kind of network presents a great number of nodes with low degree and a few nodes with high degree (Fig 4A). Nodes corresponding to topologies number one and three showed the highest degree (*deg*(1) = *deg*(3) = 14). Curiously, we found that nodes with higher degree turned out to be also those network topologies with higher abundance (Fig 4B). This relationship between abundance and node degree was corroborated by the Spearman’s rank correlation coefficient (*r_S_* = 0.353752, *p* = 6.2933 × 10^−23^).

**Fig 4.**
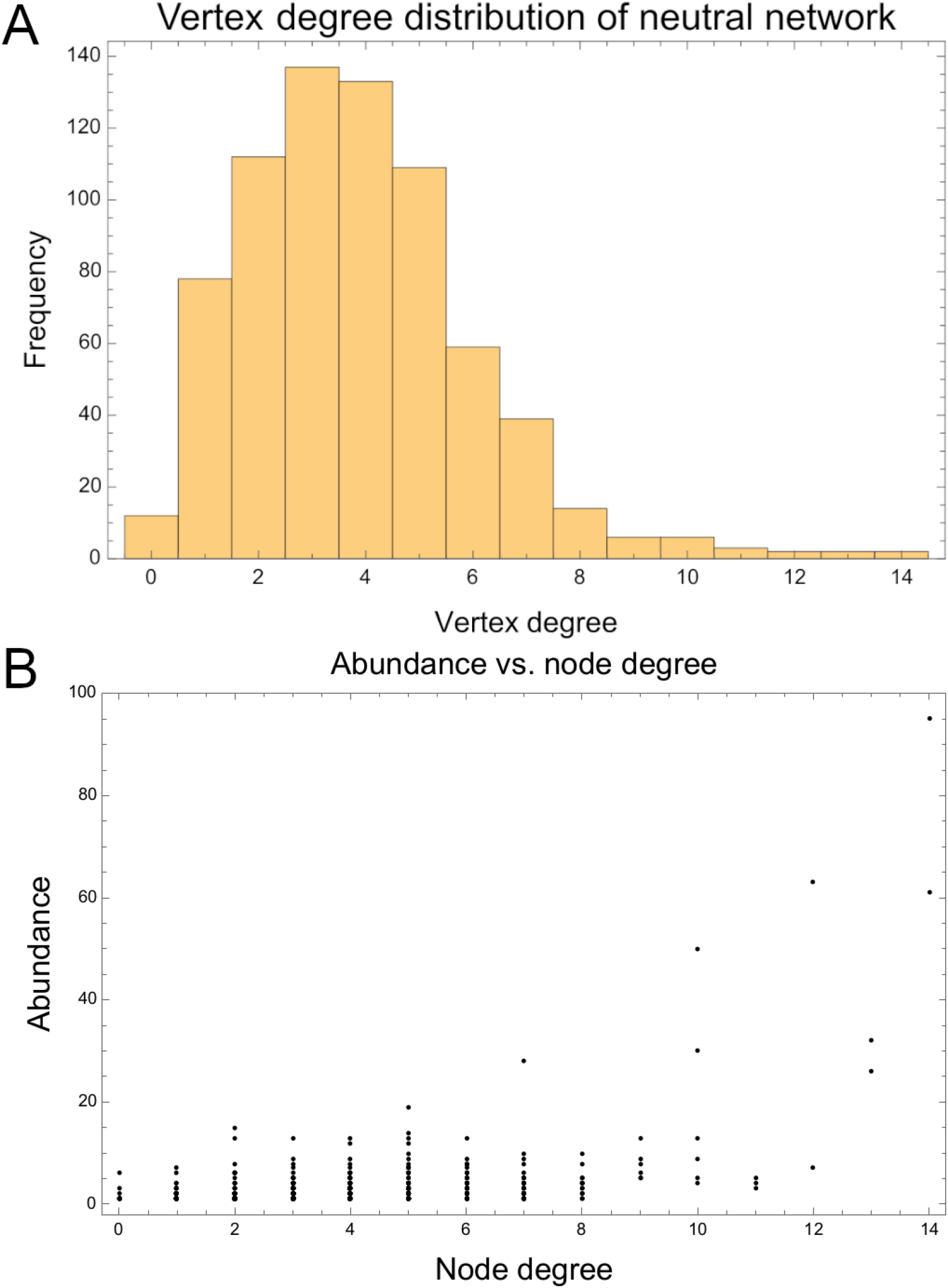
Node degree in the neutral network. (A) Histogram of vertex degree distribution in the neutral network. (B) Scatter plot of Topology abundance vs. Node degree in the neutral network.

Furthermore, we examined the extent to which the metagraph of topologies remained highly connected when joining neighbors at two mutational steps. Interestingly, we found that only topology number 420 remained isolated from the rest of topologies in the metagraph (Fig 5).

**Fig 5.**
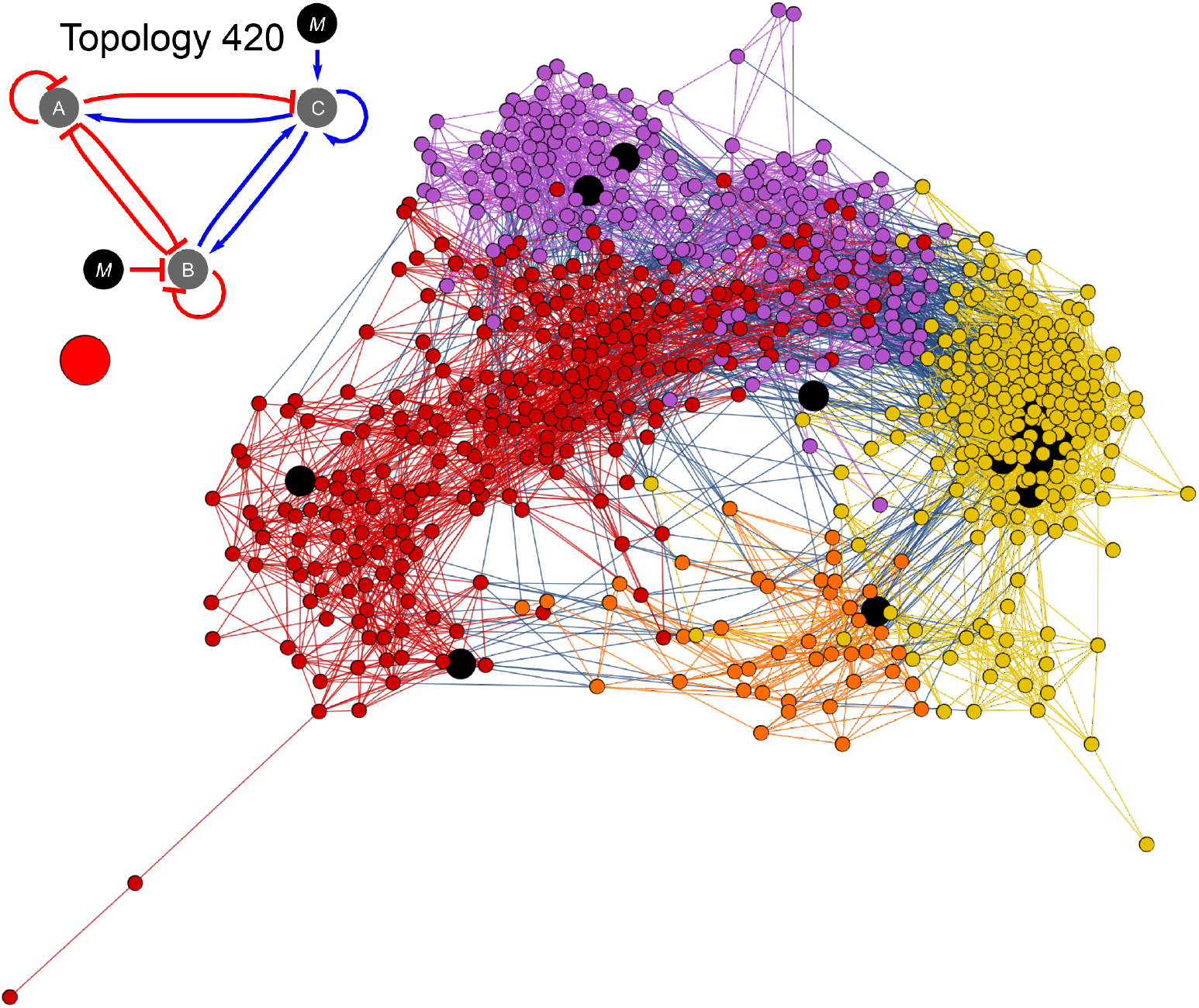
Neutral network of topologies separated by two mutational steps. The network shows that almost all the topologies are connected by two or less mutational steps. Big red circle represents topology 420 that remains unconnected from the main graph. Big black circles correspond to the 15 most abundant topologies described in Fig 3. Different colors in the graph represent different clusters of topologies.

Next to topological robustness, we investigated the robustness of the sampled network topologies to parameter changes (see Methods). Fig 6A illustrates the association between the abundance of a topology class versus its robustness. We did not find a significant correlation between these two variables (*r_S_* = −0.0389724, p-value = 0.298475). However, when we grouped those topologies containing the same subgraph and calculated their mean robustness, we found that between the most robust subgraphs is the “Bistable” motif, the I3-FFL, as well as the “Overlapping domains” motif (which is a variation of the I3-FFL), which have been previously reported in Cotterell & Sharpe [28] (see S2 Fig and S1 Table).

**Fig 6.**
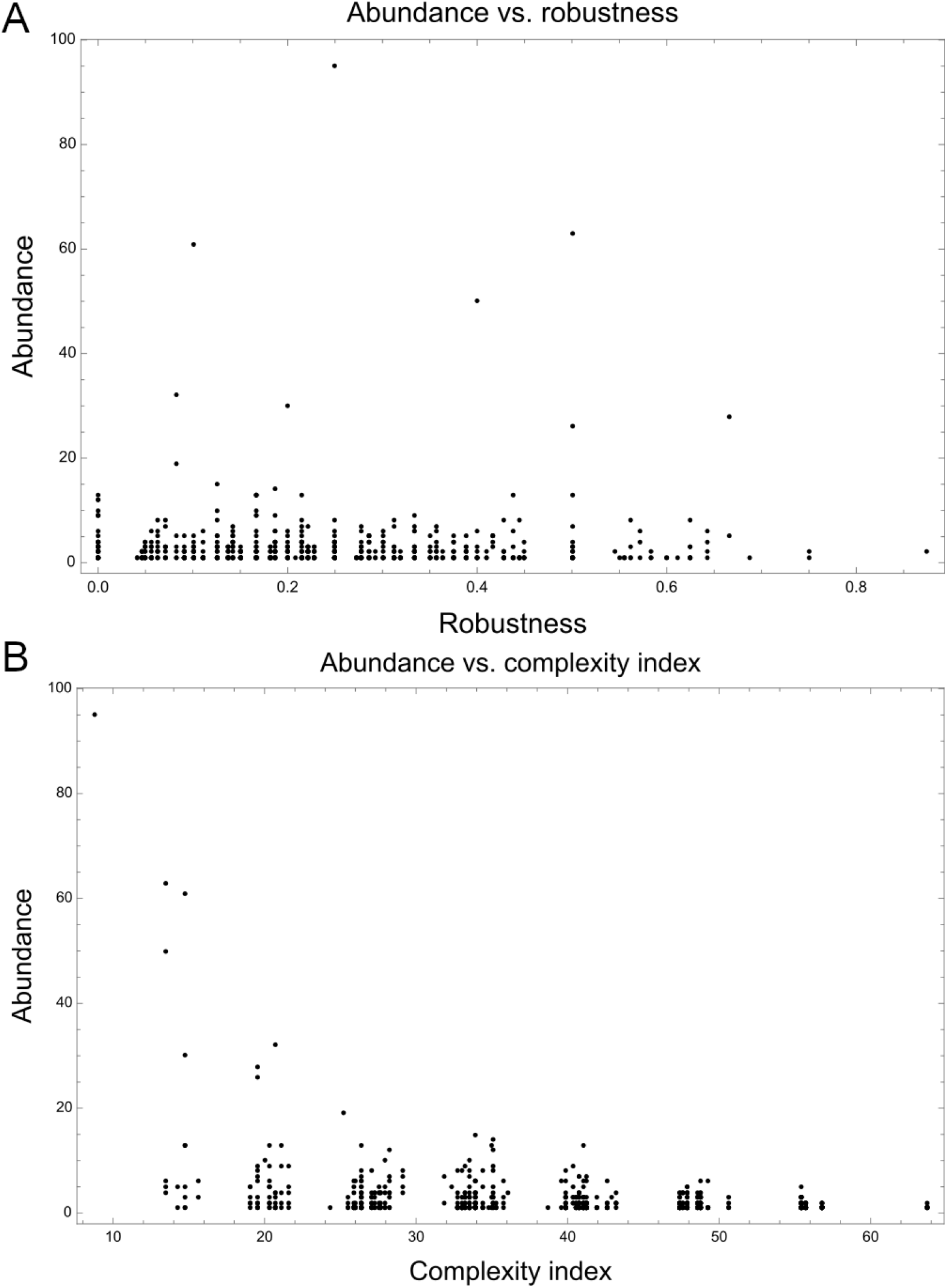
Relationship between the abundance of topologies with robustness and complexity index. (A) Abundance vs. Robustness. The robustness of each topology was calculated as the ratio of perturbations in which the fitness was greater or equal to 0.95 with respect to the total of perturbations performed. (B) Abundance vs. Complexity index. Each point in the plot represents one of the 714 network topologies. More complex topologies tend to be less abundant than simple topologies (*r_S_* = −0.316653, p-value = 2.25161 × 10^−18^).

On the other hand, analysis of the average vertex degree seems to be indicative that network topologies are likely to withstand numerous topological modifications (approximately 4) without significant loss in their ability to generate a striped pattern of gene expression, representing a high fitness solution to the prescribed task. The average complexity index of network topologies was 39.2435 (±11.6768) and was found to be negatively correlated with the abundance (*r_S_* = −0.316653, p-value = 2.25161 × 10^−18^) (Fig 6B). Additionally, the average path length seems to suggest that the evolutionary landscape could be easily traversed by most network topologies without incurring in significant fitness loss as long as mutational changes results in moves between adjacent neighbors across the metagraph (Table 1). It is interesting to note that, although the examined GRNs seem to exhibit, on average, a high robustness to topological/parameter changes, they generally show a rather high sensitivity to variation in the size of the morphogenetic field. For instance, we found that when the size of the morphogenetic field was fixed at 10, 20, 40 and 50 cells only 4.66%, 56.82%, 56.14% and 41.05%, respectively, of the examined GRNs showed a fitness greater than 0.9 (S3 FigB). In addition to robustness, we wanted to investigate which of the GRNs could still produce a striped pattern in absence of diffusion, as it has been shown that some networks can work without diffusion involved [32]. We found that out of 2061 GRNs, 404 were diffusion independent, many of them presenting the I3-FFL (see S1 Appendix).

**Table 1.** Network descriptors of neutral network of topologies.

| Network descriptor | Value |
| --- | --- |
| Average vertex degree | 3.85714 |
| Clustering coefficient [49] | 0.117097 |
| Average path length | 7.25261 |
| Graph connectedness | 0.00540974 |
| Graph diameter* | 25 |
| Average path length* | 9.0499 |

\*These descriptors were calculated for the main connected subgraph of the network.

Hierarchical clustering of spatiotemporal expression profiles showed that GRNs with topologies that seemed to be very different have indeed very similar expression dynamics. We expected to find GRNs with the same topology to group together but instead we found GRNs with different topologies grouped as close neighbors (Fig 7A). Intriguingly, some GRNs displaying very abundant network topologies were grouped together with GRNs displaying very infrequent network topologies. This seems to indicate that additional interactions in the networks are likely to represent alternative regulatory mechanisms for fine tuning gene expression, instead of drastically changing the expression mechanism, which can be seen in the expression dynamics of networks with different topologies (S4 Fig). Additionally, clustering of topologies revealed that there are six main groups of topologies in our set of 2061 GRNs (Fig 7B).

**Fig 7.**
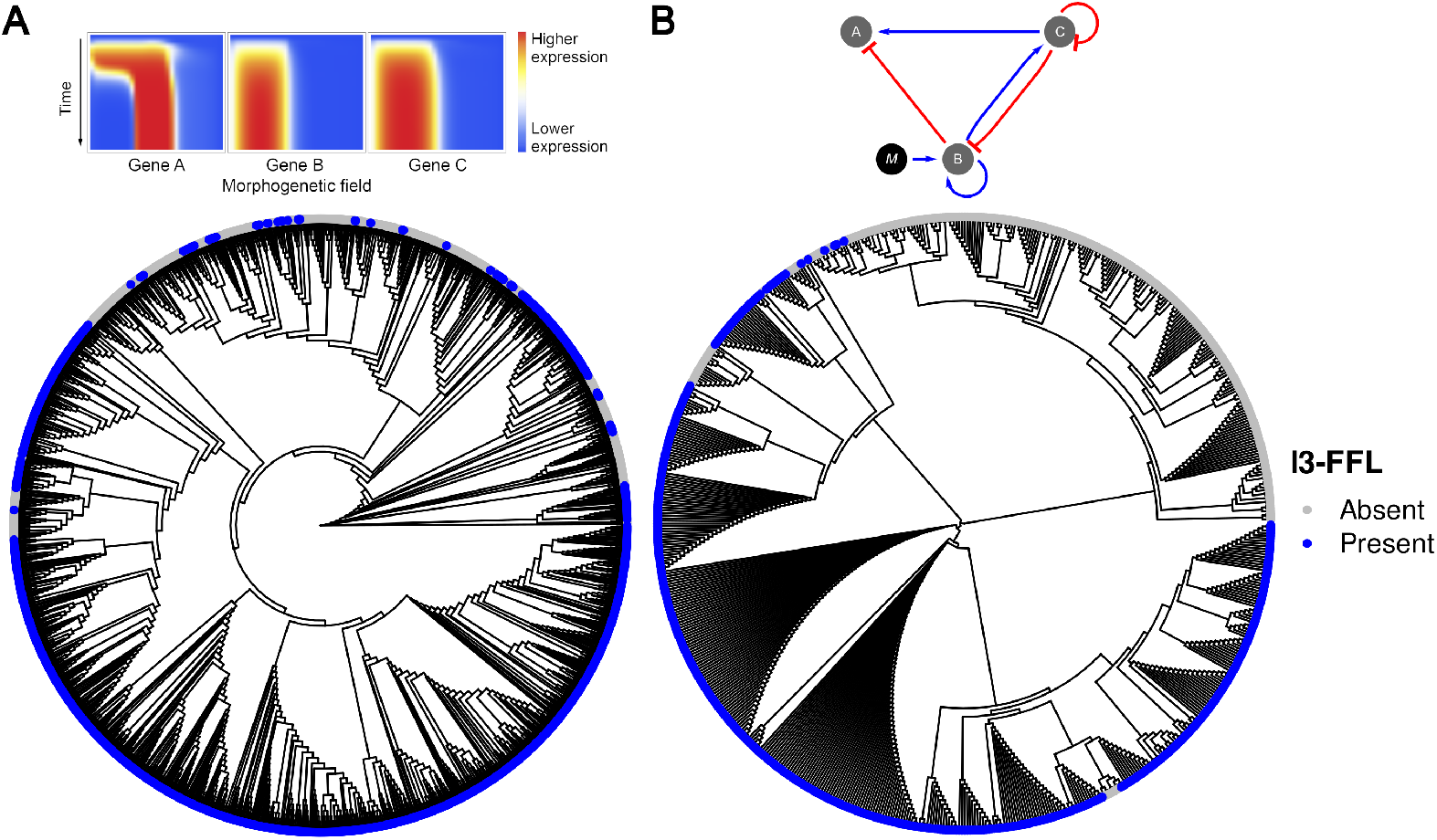
Clustering of expression profiles and topologies. (A) Clustering of spatiotemporal expression profiles obtained by neighbor joining. (B) Clustering of network topologies. Blue circles correspond to topologies that contain the I3-FFL network motif.

Among the new network topologies we report here, perhaps one of the most interesting ones is the topology number 9, which is able to express the striped gene expression pattern with only two genes (Fig 8). This topology is composed of a core quite similar to the Gierer-Meinhardt mechanism [50, 51] known to generate stripes and dots in bidimensional fields, with the only differences being the self inhibition of gene A and the inhibition of the genes by the morphogen. Such additional interactions in the topology number 9 play a role in restricting the striped expression pattern to a precise location along the morphogenetic field.

**Fig 8.**
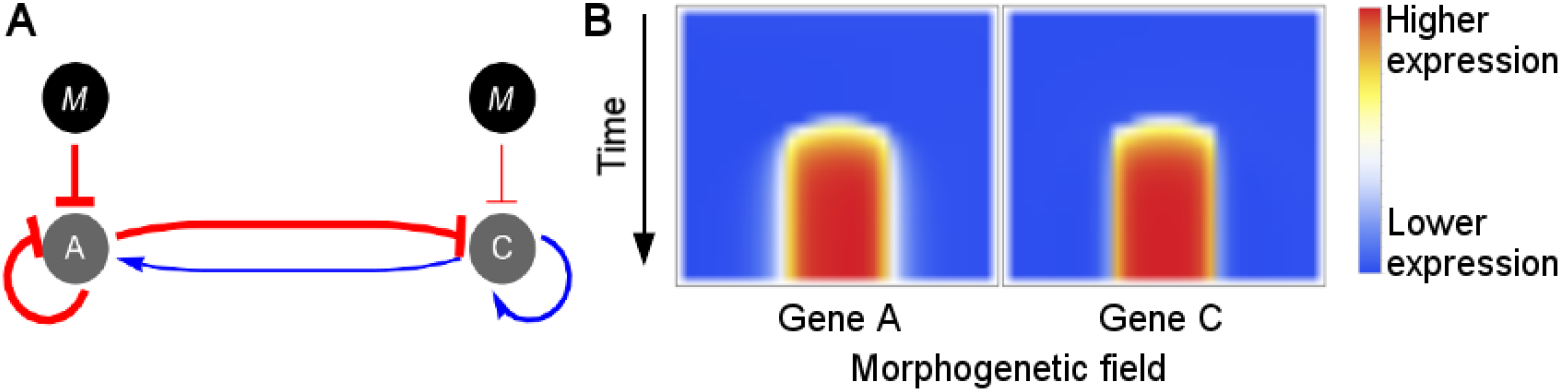
Expression dynamics of topology 9. (A) Network topology number 9. Although this topology consists only of two genes, it display a striped pattern of gene expression for both of them as can be observed in (B). (B) Spatiotemporal expression profile of a GRN with topology 9. Gene **B** is not shown as it was lost in this topology in the evolutionary process simulated by the search space algorithm.

Despite having only two genes, the regulatory mechanism underlying the topology 9 generates dynamically more complex expression patterns compared to other topologies, such as the I3-FFL; however, and despite the fact that the Gierer-Meinhardt model forms patterns that can scale with tissue size in the presence of external forcing [52, 53], we found that topology 9 was more sensible to changes in the morphogenetic field size than those topologies based on I3-FFL (S2 Table), indicating that the additional regulatory interactions need to be fine-tuned depending on the number of cells in the field.

In the mechanism of expression of topology 9 the initial expression pattern of gene A is uniform across the morphogenetic field, but later on the morphogen gradient shapes its dynamics in such a way that two important events occur: firstly, the overall expression level of the gene A is decreased by the inhibition of the morphogen, releasing gene C from its repression by gene A; next, gene A tend to display a non-homogeneous spatial distribution with higher expression levels being shifted towards the cells at the rightmost part of the morphogenetic field.

These two events lead to the repression of the gene C by the gene A in the rightmost part of the morphogenetic field and by the morphogen in the leftmost part of the morphogenetic field, thus generating a slightly higher level of expression of gene C in the center of the field. Because gene C activates its own expression, a band of expression begins to form. This band of expression then begins to activate the gene A also in the center of the field (S5 Fig).

In this way two bands of expression that depend on each other are formed. The high level of gene C in the middle of the field counteract its inhibition by gene A, while at the ends of the bands the inhibition of gene A on gene C overcomes the self-activation of gene C, preventing the expression of the latter to extend out of the middle of the field. In turn, the band of expression of gene A can only exist in the middle of the field because its expression depends on gene C.

This is a complex mechanism reminiscent of the Gierer-Meinhardt model, which to the best of our knowledge is reported for the first time in the context of multiple morphogen inputs, and could thus be of particular interest for synthetic biology implementations to assess the feasibility and robustness conferred by this mechanism to GRNs.

## Discussion

In this work, we have conducted extensive computational explorations of small GRNs with the capacity to generate a simple spatial pattern in a 1-D morphogenetic field. Our findings shed new light on regulatory rules and dynamical mechanisms that GRNs in nature could potentially implement to achieve stereotyped spatio-temporal tasks.

Overall, based on our extensive search space exploration we identified a great variety of pattern-forming GRN topologies with distinctive and often widely shared building blocks (network motifs), which are likely to confer each topology the ability to achieve specific regulatory tasks in a semi-autonomous manner. In particular, we uncovered a great variety of network topologies that could produce a striped gene expression pattern, where 615 out of 714 were found to be multi-input topologies, which highlights the importance of often disregarded pleiotropic signaling events as a potentially critical design specification rule for successful synthetic biology implementations of GRNs with various types of pattern-forming abilities. Most importantly, we found that this ensemble of pattern-forming GRNs tends to form a highly connected meta-graph, which highlights emergent properties, such as robustness and evolvability, of complex biological systems at a very high level [54].

Robustness is a defining feature of any developmental patterning process. Our results suggest that the underlying morphogen interpretation mechanisms (GRNs) are highly accessible by exploring the topology space constrained by selection for their final steady-state and by their ability to buffer noise [37, 55]. In our study, the most abundant topologies were found forming highly connected clusters in a so-called neutral network of genotypes, implying that the corresponding GRN topologies would tend to be robust to quantitative changes in the strength of regulatory interactions, diffusion, and degradation parameters, as well as to qualitative changes in the topology itself. In essence, such organization could be seen as an intrinsic evolutionary property of GRNs that facilitates the coexistence of robustness and evolvability [56]. For instance, bridge topologies in the neutral network connect distinct classes of pattern-forming GRNs (S6 Fig), suggesting that evolutionary transitions between different GRNs (for example, between a Gierer-Meinhardt-like network to an Incoherent Feed-Forward Loop) could be achieved with minimal fitness costs. This high-level property of the neutral network of topologies could thus facilitate the fine-tuning of the regulatory systems through genetic modifications to meet specific functional requirements under selective pressures.

In contrast to this robustness to qualitative and quantitative changes, we found that many GRNs were susceptible to changes in morphogenetic field size. However, GRNs with the same topology behave in different ways for each field size (S3 FigC), indicating that the specific values of network parameters are more important than the network topology when it comes to producing the same pattern at different scales.

Other studies have shown that Incoherent Feed-Forward Loops can form spatial stripes of gene expression [57]. In particular, we found that the I3-FFL is one of the network motifs that appears more frequently in these GRNs, which is in agreement with the study by Cotterell & Sharpe [28], and has been found in transcriptional networks involved in cell fate definition [58]. Moreover, our results indicate that this is a robust motif that can be useful for constructing synthetic gene regulatory networks exposed to stochastic fluctuations in the environment and mutations.

Moreover, as our model was less stringent and allowed the morphogen input to interact with any gene, we found a significantly larger number of pattern-forming topologies, which can operate under disparate combinations of diffusion and degradation parameters (S7 Fig). For example, in a single input GRN model, the I4-FFL has not been found capable of generating a striped gene expression pattern [27], whereas, in our experimental setup, 20.17% of the topologies were found to implement such a regulatory motif (S1 Table).

Although recent experimental evidence highlights the importance of the Incoherent Feed-Forward Loop type 2 (I2-FFL) for generating a striped gene expression pattern [16, 35], it is intriguing to find that such motif is underrepresented in our study, with only 10.92% of the topologies implementing such regulatory mechanism. This might be due to the difficulty to find optimal solutions when probing the design space of multi-input GRNs, which are likely to contain multiple locally optimal solutions. In addition, it is important to mention here that the application of a MCMC-like algorithm as a search space strategy for such complex tasks cannot be expected to allow for extensive exploration of the design space of GRNs nor for the accessibility of every single locally optimal solution. However, it is also possible that the I2-FFL motif is not as robust as other motifs and therefore be less represented.

We also found GRNs with a similar topology to that of the “opposing gradients” reported by Schaerli et al. [32, 35]. Although the “opposing gradients” mechanism requires the constitutive expression of two of the genes in the GRN, and our model does not take constitutive expression into account, we observed that the GRNs that implement the “opposing gradients” mechanism also tend to implement auto-regulation interactions on these two nodes, bypassing in this way the lack of constitutive expression as found in previous studies [27].

The correlation between the complexity index of a topology and its abundance indicates that networks with simpler designs are more robust to changes in interaction parameters and that more complex networks are less robust. For example, topology number 420 (Fig 5), which presents high complexity, was one of the topologies with lower abundance and was disconnected from all other topologies in the two-step neighbors in the neutral network.

Although GRNs that produce a band of gene expression with only two nodes have been described (e.g., the I-zero motif [35]), this is to our knowledge the first time that a novel GRN topology such as number 9 (Fig 8) has been reported, which adds considerably to existing knowledge on the genotype-phenotype mapping problems studied in the context of developmental pattern formation. It would be interesting to assess experimentally the ability of this particular topology to generate striped gene expression patterns, as well as to assess its robustness in the face of mutational changes and noisy morphogen input profiles.

Overall, we believe our work provides a set of enticing hypotheses on GRN designs, represented as a large catalog of distinct GRN topologies that could provide valuable information as starting points for future computational and theoretical analysis of the genotype-phenotype map, as well as for experimental validation and future discovery of potentially interesting GRN designs.

## Supporting information

**S1 Fig. Most GRNs reach the steady state before 250 time steps.** (A) Gene regulatory network displaying the topology number 33. (B) Spatiotemporal expression profile of the gene regulatory network shown in (A).

**S2 Fig. Relevant subgraphs present in the set of GRNs.** These are subgraphs that have been reported in previous studies as networks involved in morphogenesis and development [28, 32, 35]. These were used to calculate the subgraph profile and the results reported in S1 Table.

**S3 Fig. Response of GRNs to changes in morphogenetic field size.** (A) Morphogen gradients of fields with 10, 20, 40 and 50 cells. These morphogen gradients were set up so that the morphogen concentration in the first cell and in the last cell were constant and only one morphogen gradient was considered at a time depending on the number of cells in the morphogenetic field. (B) Fitness of the GRNs by topology in each one of the morphogenetic fields. (C) Principal Component Analysis of fitness by morphogenetic field size. Although principal components separate GRNs in clusters, not all the GRNs with the same topology are located in the same group. (D) Average fitness of GRNs evolved in different morphogenetic field sizes.

**S4 Fig. Expression dynamics of main networks.** The dynamics of expression is shown from *t* = 1 to *t* = 30 for the eight most abundant topologies. The red line represents the expression level of the gene C along the morphogenetic field, the blue line represents the expression level of gene B and the dotted line represents the expression level of gene A.

**S5 Fig. Dynamics of expression of topology number 9.** The dynamics of expression is shown from *t* = 5 to *t* = 150, where this topology reaches its steady state expression. The red line represents the expression level of the gene C along the morphogenetic field, whereas the dotted line represents the expression level of gene A. The striped pattern of gene expression can be seen for both genes since *t* = 80.

**S6 Fig. Bridge topologies in the neutral network.** “Bridge topologies” are those topologies that connect two clusters in the neutral network (Fig 3), and as such can be interpreted as intermediary steps in the evolution from a cluster of related topologies into another cluster. For example the topology 441 could be an initial step to reach the topology 9, as in this topology nodes B and C are not connected.

**S7 Fig. Diffusion and degradation parameters.** (A) Ternary plot showing the combination of diffusion parameters for genes A, B and C. (B) Ternary plot showing the combination of degradation parameters for genes A, B and C. The color gradient that goes from topology 1 in white to topology 714 in black shows no distinguishable pattern.

**S1 Appendix. Diffusion independent networks and robustness to changes in the signal input.**

**S1 Table. Proportion of relevant subgraphs in the set of GRNs.** The 19 subgraphs presented in this table are those presented in S2 Fig. Most of the topologies presented at least one negative feedback loop (70.73%), and 62.46% of them presented the I3-FFL network motif. The robustness of each subgraph was calculated as the mean robustness of the topologies presenting that subgraph.

**S2 Table. Average fitness of each topology in different morphogenetic fields.** The average fitness by topology is reported for morphogenetic fields with 10, 20, 40 and 50 cells. Data from this table are shown in S3 FigB.

## Acknowledgements

The computational resources (Stevin Supercomputer Infrastructure) and services used in this work were provided by the VSC (Flemish Supercomputer Center), funded by Ghent University, FWO and the Flemish Government–department EWI. All authors acknowledge partial support from CODI-Universidad de Antioquia under project “Estudio de la Evolución de Circuitos de Regulación del Desarollo Embrionario a través de la Biología Evolutiva de Sistemas” code 2017-14367. All authors acknowledge the Colombian youth for their defense of a better country.

## References

1. Turing AM. The chemical basis of morphogenesis. Bulletin of Mathematical Biology. 1952;52(1-2):153–197. doi:10.1007/BF02459572.

2. Wolpert L. Positional information and the spatial pattern of cellular differentiation. Journal of Theoretical Biology. 1969;25(1):1–47. doi:https://doi.org/10.1016/S0022-5193(69)80016-0.

3. Sharpe J. Wolpert’s French Flag: what’s the problem? Development (Cambridge, England). 2019;146(24). doi:10.1242/dev.185967.

4. Driever W, Nüsslein-Volhard C. A gradient of bicoid protein in Drosophila embryos. Cell. 1988;54(1):83–93. doi:10.1016/0092-8674(88)90182-1.

5. Driever W, Nüsslein-Volhard C. The bicoid protein determines position in the Drosophila embryo in a concentration-dependent manner. Cell. 1988;54(1):95–104. doi:10.1016/0092-8674(88)90183-3.

6. Affolter M, Basler K. The Decapentaplegic morphogen gradient: from pattern formation to growth regulation. Nature Reviews Genetics. 2007;8(9):663–674. doi:10.1038/nrg2166.

7. Dessaud E, Ribes V, Balaskas N, Yang LL, Pierani A, Kicheva A, et al. Dynamic Assignment and Maintenance of Positional Identity in the Ventral Neural Tube by the Morphogen Sonic Hedgehog. PLOS Biology. 2010;8(6):e1000382. doi:10.1371/journal.pbio.1000382.

8. Cohen M, Page KM, Perez-Carrasco R, Barnes CP, Briscoe J. A theoretical framework for the regulation of Shh morphogen-controlled gene expression. Development. 2014;141(20):3868–3878. doi:10.1242/dev.112573.

9. Raspopovic J, Marcon L, Russo L, Sharpe J. Digit patterning is controlled by a Bmp-Sox9-Wnt Turing network modulated by morphogen gradients. Science. 2014;345(6196):566. doi:10.1126/science.1252960.

10. Milo R, Shen-Orr SS, Itzkovitz S, Kashtan N, Chklovskii D, Alon U. Network Motifs: Simple Building Blocks of Complex Networks. Science. 2002;298(5594):824–827. doi:10.1126/science.298.5594.824.

11. Kitano H. Computational systems biology. Nature. 2002;420(6912):206–210. doi:10.1038/nature01254.

12. Kalir S, Mangan S, Alon U. A coherent feed-forward loop with a SUM input function prolongs flagella expression in Escherichia coli; 2005.

13. Kalir S, Alon U. Using a quantitative blueprint to reprogram the dynamics of the flagella gene network. Cell. 2004;117(6):713–720. doi:10.1016/j.cell.2004.05.010.

14. Mangan S, Zaslaver A, Alon U. The coherent feedforward loop serves as a sign-sensitive delay element in transcription networks. Journal of molecular biology. 2003;334(2):197–204.

15. O’Donnell KA, Wentzel EA, Zeller KI, Dang CV, Mendell JT. c-Myc-regulated microRNAs modulate E2F1 expression. Nature. 2005;435(7043):839–843. doi:10.1038/nature03677.

16. Basu S, Gerchman Y, Collins CH, Arnold FH, Weiss R. A synthetic multicellular system for programmed pattern formation. Nature. 2005;434(7037):1130–1134. doi:10.1038/nature03461.

17. Atkinson MR, Savageau MA, Myers JT, Ninfa AJ. Development of genetic circuitry exhibiting toggle switch or oscillatory behavior in Escherichia coli. Cell. 2003;113(5):597–607. doi:10.1016/s0092-8674(03)00346-5.

18. Friedland AE, Lu TK, Wang X, Shi D, Church G, Collins JJ. Synthetic gene networks that count. Science (New York, NY). 2009;324(5931):1199–1202. doi:10.1126/science.1172005.

19. Gardner L, Deiters A. Light-controlled synthetic gene circuits. Current Opinion in Chemical Biology. 2012;16(3-4):292–299. doi:10.1016/j.cbpa.2012.04.010.

20. Levskaya A, Chevalier AA, Tabor JJ, Simpson ZB, Lavery LA, Levy M, et al. Engineering Escherichia coli to see light. Nature. 2005;438(7067):441–442. doi:10.1038/nature04405.

21. Santos-Moreno J, Schaerli Y. Using Synthetic Biology to Engineer Spatial Patterns. Advanced Biosystems. 2019;3(4):1800280. doi:https://doi.org/10.1002/adbi.201800280.

22. Karlsson M, Weber W. Therapeutic synthetic gene networks. Current Opinion in Biotechnology. 2012;23(5):703–711. doi:10.1016/j.copbio.2012.01.003.

23. Higashikuni Y, Chen WC, Lu TK. Advancing therapeutic applications of synthetic gene circuits. Current Opinion in Biotechnology. 2017;47:133–141. doi:10.1016/j.copbio.2017.06.011.

24. Abil Z, Xiong X, Zhao H. Synthetic biology for therapeutic applications. Molecular Pharmaceutics. 2015;12(2):322–331. doi:10.1021/mp500392q.

25. Healy CP, Deans TL. Genetic circuits to engineer tissues with alternative functions. Journal of Biological Engineering. 2019;13:39. doi:10.1186/s13036-019-0170-7.

26. Kitada T, DiAndreth B, Teague B, Weiss R. Programming gene and engineered-cell therapies with synthetic biology. Science (New York, NY). 2018;359(6376). doi:10.1126/science.aad1067.

27. Munteanu A, Cotterell J, Solé RV, Sharpe J. Design principles of stripe-forming motifs: the role of positive feedback. Scientific Reports. 2014;4(1):5003. doi:10.1038/srep05003.

28. Cotterell J, Sharpe J. An atlas of gene regulatory networks reveals multiple three-gene mechanisms for interpreting morphogen gradients. Molecular Systems Biology. 2010;6(425):1–14. doi:10.1038/msb.2010.74.

29. von Dassow G, Meir E, Munro EM, Odell GM. The segment polarity network is a robust developmental module. Nature. 2000;406(6792):188–192. doi:10.1038/35018085.

30. Reinitz J, Mjolsness E, Sharp DH. Model for cooperative control of positional information in Drosophila by bicoid and maternal hunchback. Journal of Experimental Zoology. 1995;271(1):47–56. doi:10.1002/jez.1402710106.

31. Jaeger J, Blagov M, Kosman D, Kozlov KN, Manu, Myasnikova E, et al. Dynamical analysis of regulatory interactions in the gap gene system of Drosophila melanogaster. Genetics. 2004;167(4):1721–1737. doi:10.1534/genetics.104.027334.

32. Schaerli Y, Jiménez A, Duarte JM, Mihajlovic L, Renggli J, Isalan M, et al. Synthetic circuits reveal how mechanisms of gene regulatory networks constrain evolution. Molecular Systems Biology. 2018;14(9):e8102. doi:10.15252/msb.20178102.

33. Elowitz MB, Leibier S. A synthetic oscillatory network of transcriptional regulators. Nature. 2000;403(6767):335–338. doi:10.1038/35002125.

34. Barbier I, Perez-Carrasco R, Schaerli Y. Controlling spatiotemporal pattern formation in a concentration gradient with a synthetic toggle switch. Molecular Systems Biology. 2020;16(6):e9361. doi:10.15252/msb.20199361.

35. Schaerli Y, Munteanu A, Gili M, Cotterell J, Sharpe J, Isalan M. A unified design space of synthetic stripe-forming networks. Nature Communications. 2014;5(May):4905. doi:10.1038/ncomms5905.

36. Balaskas N, Ribeiro A, Panovska J, Dessaud E, Sasai N, Page KM, et al. Gene Regulatory Logic for Reading the Sonic Hedgehog Signaling Gradient in the Vertebrate Neural Tube. Cell. 2012;148(1):273–284. doi:10.1016/j.cell.2011.10.047.

37. Exelby K, Herrera-Delgado E, Perez LG, Perez-Carrasco R, Sagner A, Metzis V, et al. Precision of tissue patterning is controlled by dynamical properties of gene regulatory networks. Development. 2021;148(dev197566). doi:10.1242/dev.197566.

38. Ricci F, Vallée-Bélisle A, Plaxco KW. High-precision, in vitro validation of the sequestration mechanism for generating ultrasensitive dose-response curves in regulatory networks. PLoS computational biology. 2011;7(10):e1002171. doi:10.1371/journal.pcbi.1002171.

39. Mendoza L, Xenarios I. A method for the generation of standardized qualitative dynamical systems of regulatory networks. Theoretical Biology & Medical Modelling. 2006;3:13. doi:10.1186/1742-4682-3-13.

40. Vu TT, Vohradsky J. Nonlinear differential equation model for quantification of transcriptional regulation applied to microarray data of Saccharomyces cerevisiae. Nucleic Acids Research. 2007;35(1):279–287. doi:10.1093/nar/gkl1001.

41. Zhang Q, Bhattacharya S, Andersen ME. Ultrasensitive response motifs: basic amplifiers in molecular signalling networks. Open Biology. 2013;3(4):130031. doi:10.1098/rsob.130031.

42. Armao JJ, Lehn JM. Nonlinear Kinetic Behavior in Constitutional Dynamic Reaction Networks. Journal of the American Chemical Society. 2016;138(51):16809–16814. doi:10.1021/jacs.6b11107.

43. Chen KC, Wang TY, Tseng HH, Huang CYF, Kao CY. A stochastic differential equation model for quantifying transcriptional regulatory network in Saccharomyces cerevisiae. Bioinformatics (Oxford, England). 2005;21(12):2883–2890. doi:10.1093/bioinformatics/bti415.

44. Dahlquist KD, Fitzpatrick BG, Camacho ET, Entzminger SD, Wanner NC. Parameter Estimation for Gene Regulatory Networks from Microarray Data: Cold Shock Response in Saccharomyces cerevisiae. Bulletin of Mathematical Biology. 2015;77(8):1457–1492. doi:10.1007/s11538-015-0092-6.

45. scikit-bio development team T. scikit-bio: A Bioinformatics Library for Data Scientists, Students, and Developers; 2020. Available from: http://scikit-bio.org.

46. Bonchev DG, Rouvray DH. Complexity in Chemistry, Biology, and Ecology, ser. Mathematical and Computational Chemistry. Springer; 2005.

47. Pedregosa F, Varoquaux G, Gramfort A, Michel V, Thirion B, Grisel O, et al. Scikit-learn: Machine Learning in Python. Journal of Machine Learning Research. 2011;12:2825–2830.

48. Erdös P, Rényi a. On random graphs. Publicationes Mathematicae. 1959;6:290–297. doi:10.2307/1999405.

49. Watts DJ, Strogatz SH. Collective dynamics of ‘small-world’ networks. Nature. 1998;393(6684):440–2. doi:10.1038/30918.

50. Gierer A, Meinhardt H. A theory of biological pattern formation. Kybernetik. 1972;12(1):30–39. doi:10.1007/BF00289234.

51. Meinhardt H, Gierer A. Pattern formation by local self-activation and lateral inhibition. BioEssays: News and Reviews in Molecular, Cellular and Developmental Biology. 2000;22(8):753–760. doi:10.1002/1521-1878(200008)22:8¡753::AID-BIES9¿3.0.CO;2-Z.

52. Shoaf SA, Conway K, Hunt RK. Application of reaction-diffusion models to cell patterning in Xenopus retina. Initiation of patterns and their biological stability. Journal of Theoretical Biology. 1984;109(3):299–329. doi:10.1016/s0022-5193(84)80085-5.

53. Ishihara S, Kaneko K. Turing pattern with proportion preservation. Journal of Theoretical Biology. 2006;238(3):683–693. doi:10.1016/j.jtbi.2005.06.016.

54. Kitano H. Biological robustness. Nature Reviews Genetics. 2004;5(11):826–837. doi:10.1038/nrg1471.

55. Lo WC, Zhou S, Wan FYM, Lander AD, Nie Q. Robust and precise morphogen-mediated patterning: trade-offs, constraints and mechanisms. Journal of The Royal Society Interface. 2015;12(102):20141041. doi:10.1098/rsif.2014.1041.

56. Wagner A. Robustness and evolvability: a paradox resolved. Proceedings of the Royal Society B: Biological Sciences. 2008;275(1630):91–100. doi:10.1098/rspb.2007.1137.

57. Ishihara S, Fujimoto K, Shibata T. Cross talking of network motifs in gene regulation that generates temporal pulses and spatial stripes. Genes to Cells. 2005;10(11):1025–1038. doi:https://doi.org/10.1111/j.1365-2443.2005.00897.x.

58. Li L, Rispoli R, Patient R, Ciau-Uitz A, Porcher C. Etv6 activates vegfa expression through positive and negative transcriptional regulatory networks in Xenopus embryos. Nature Communications. 2019;10(1). doi:10.1038/s41467-019-09050-y.

