## Supplementary figures and images for "Elucidating multi-input processing 3-node gene regulatory network topologies capable of generating striped gene expression patterns"

### S1 Fig

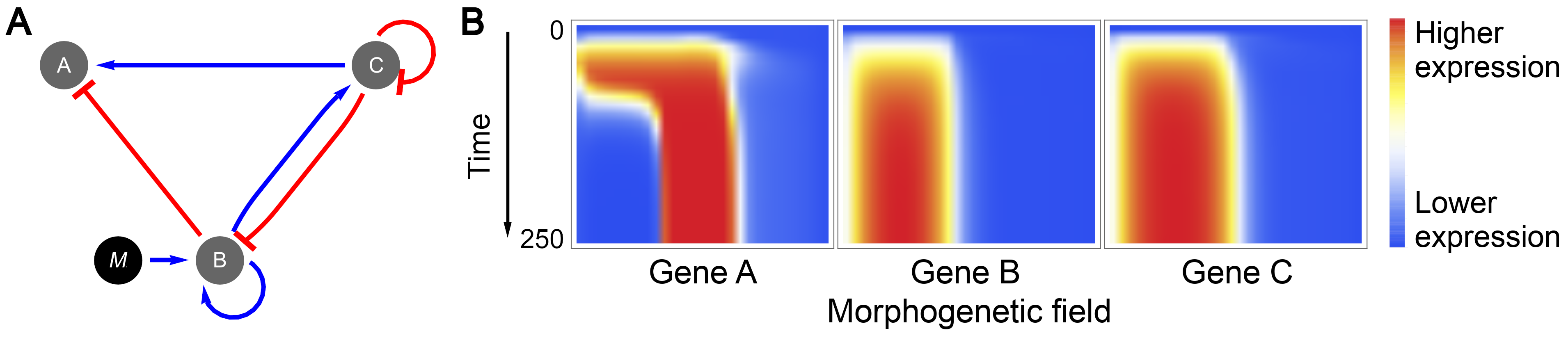

### S2 Fig

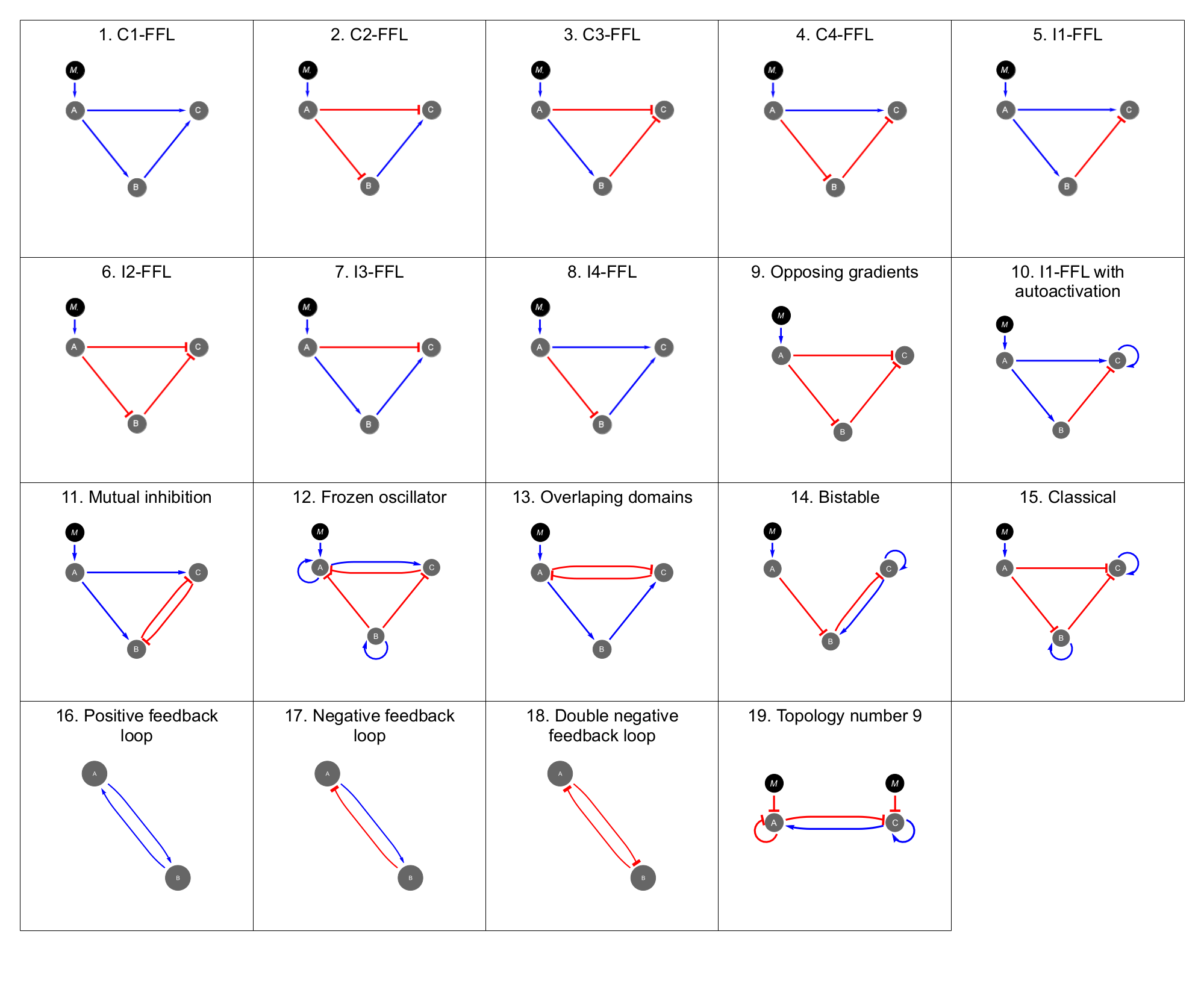

### S3 Fig

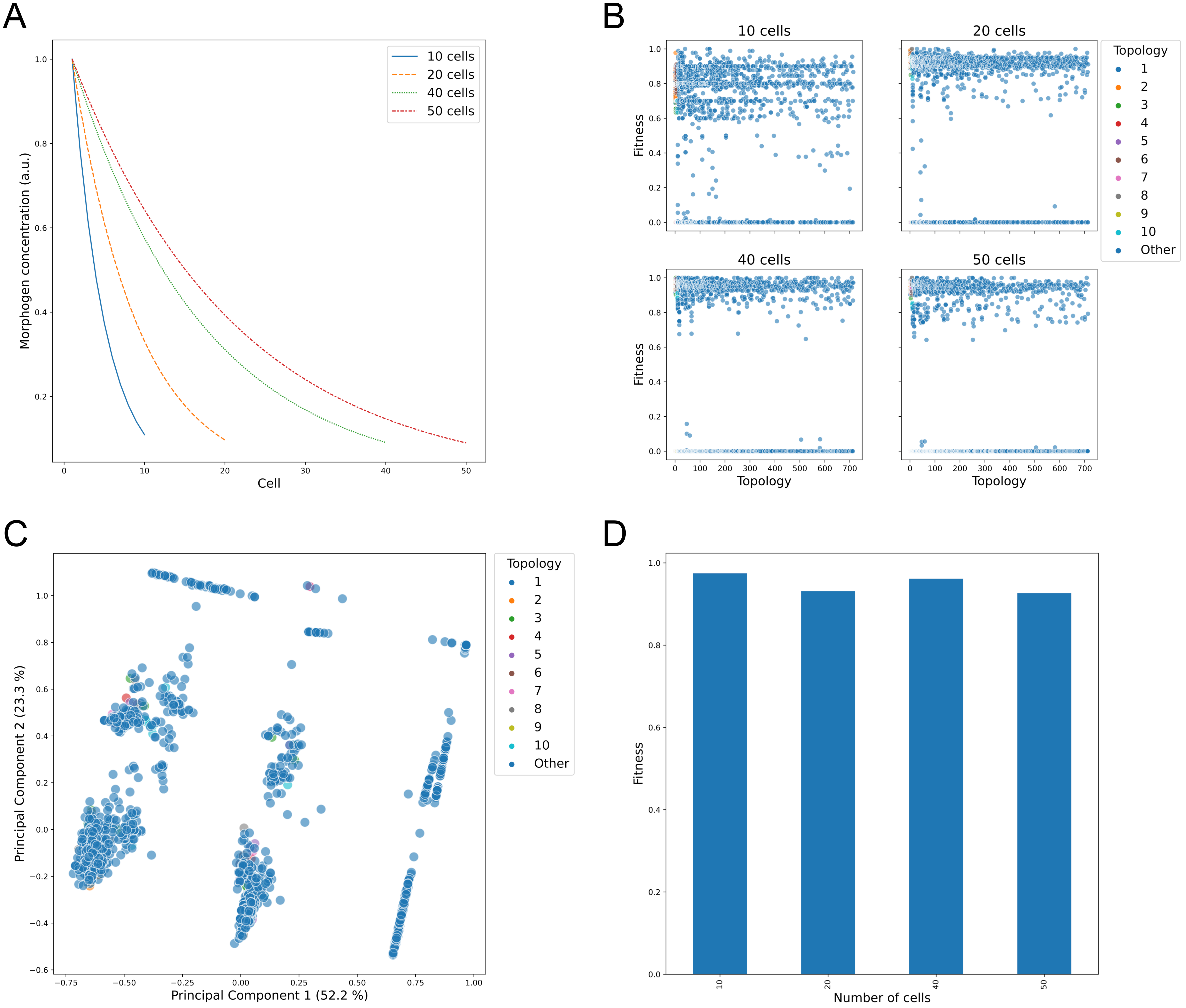

### S4 Fig

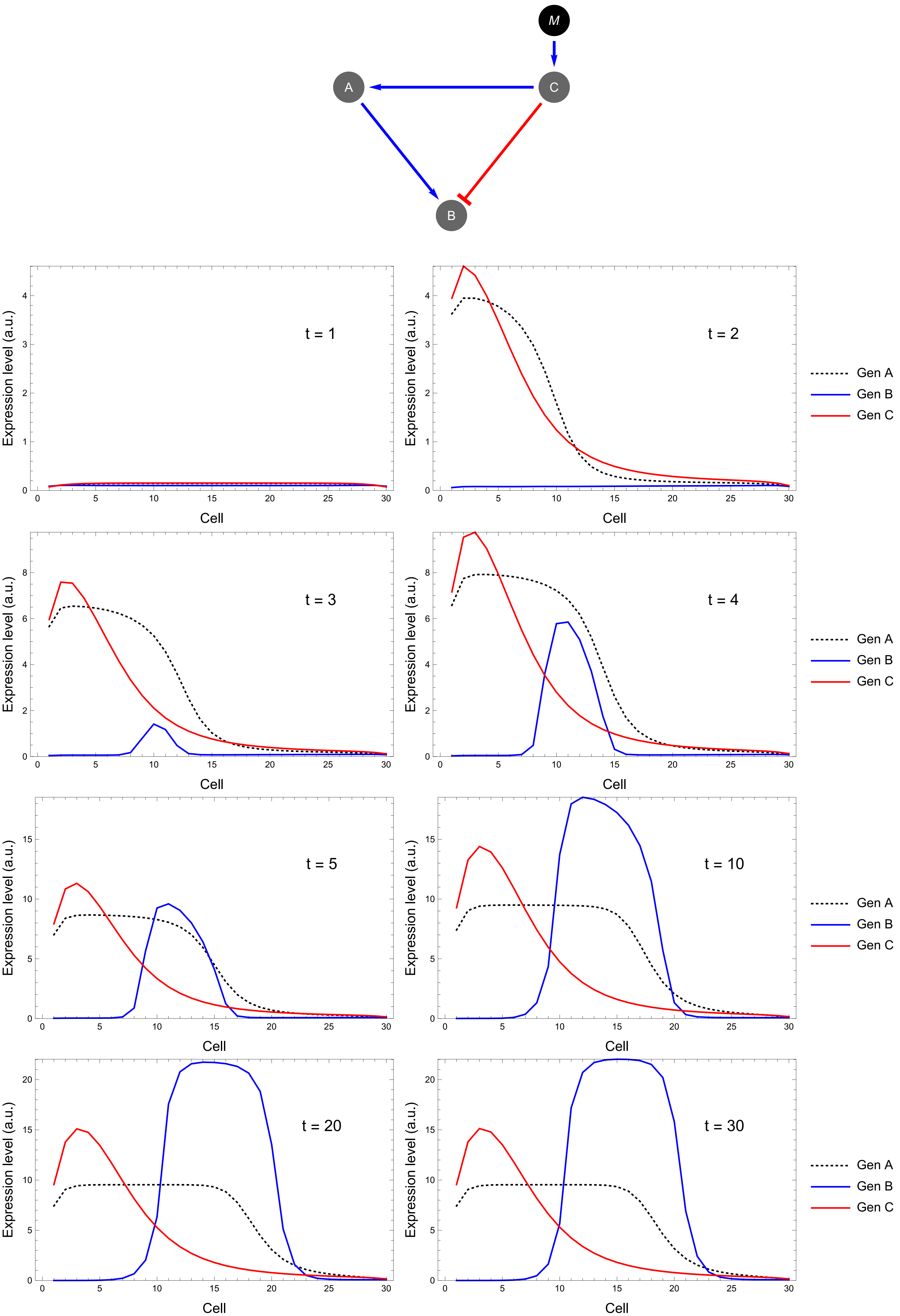

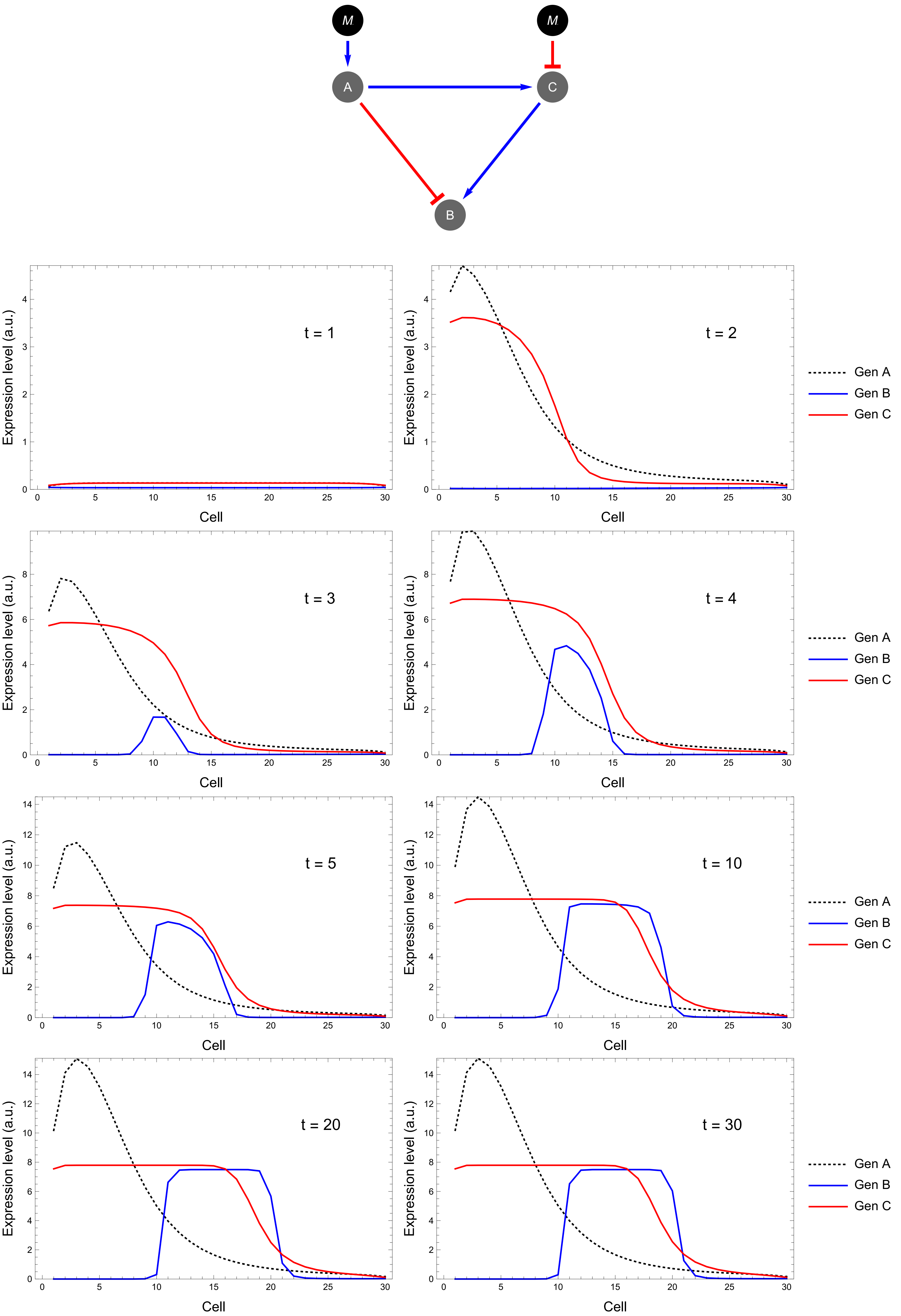

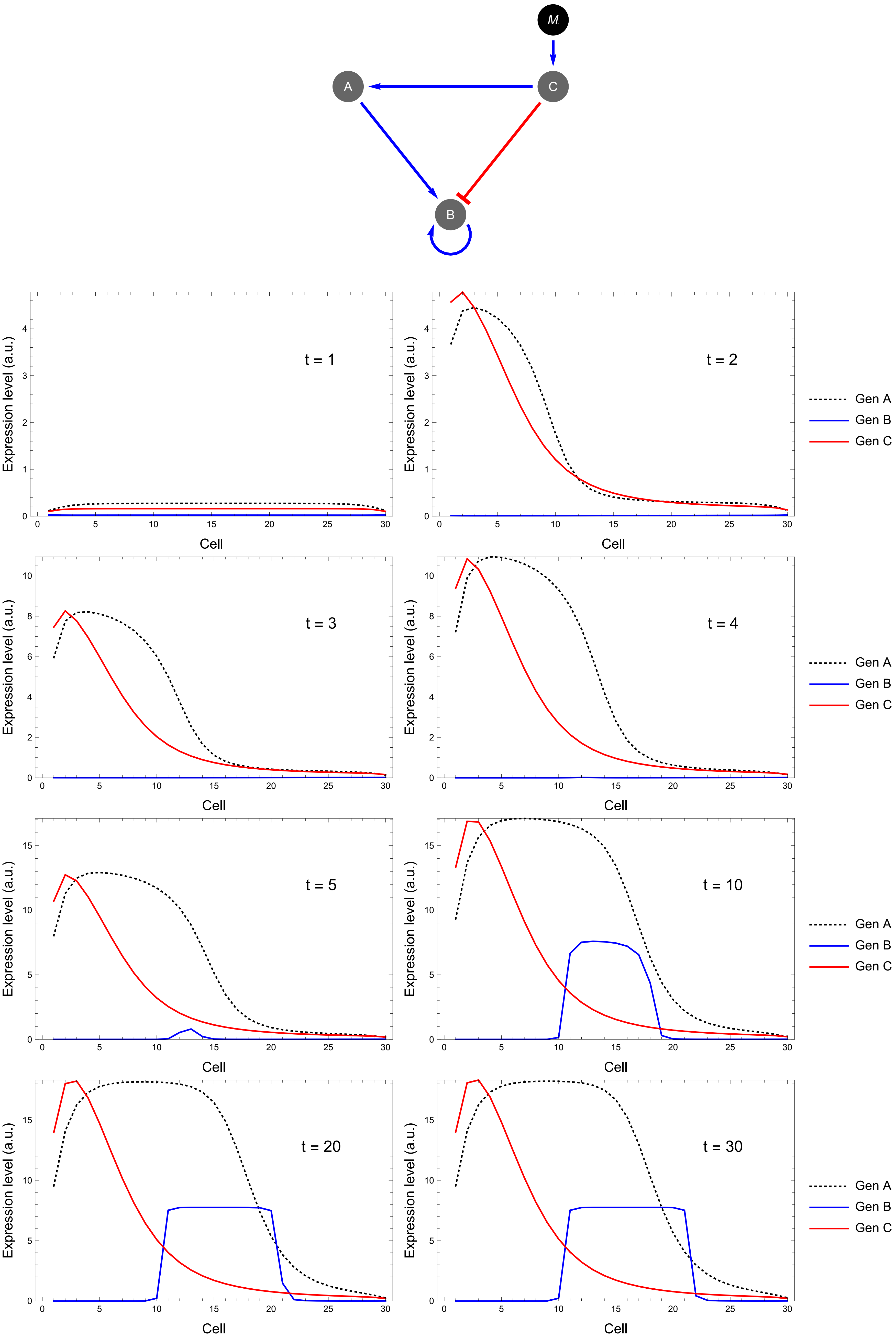

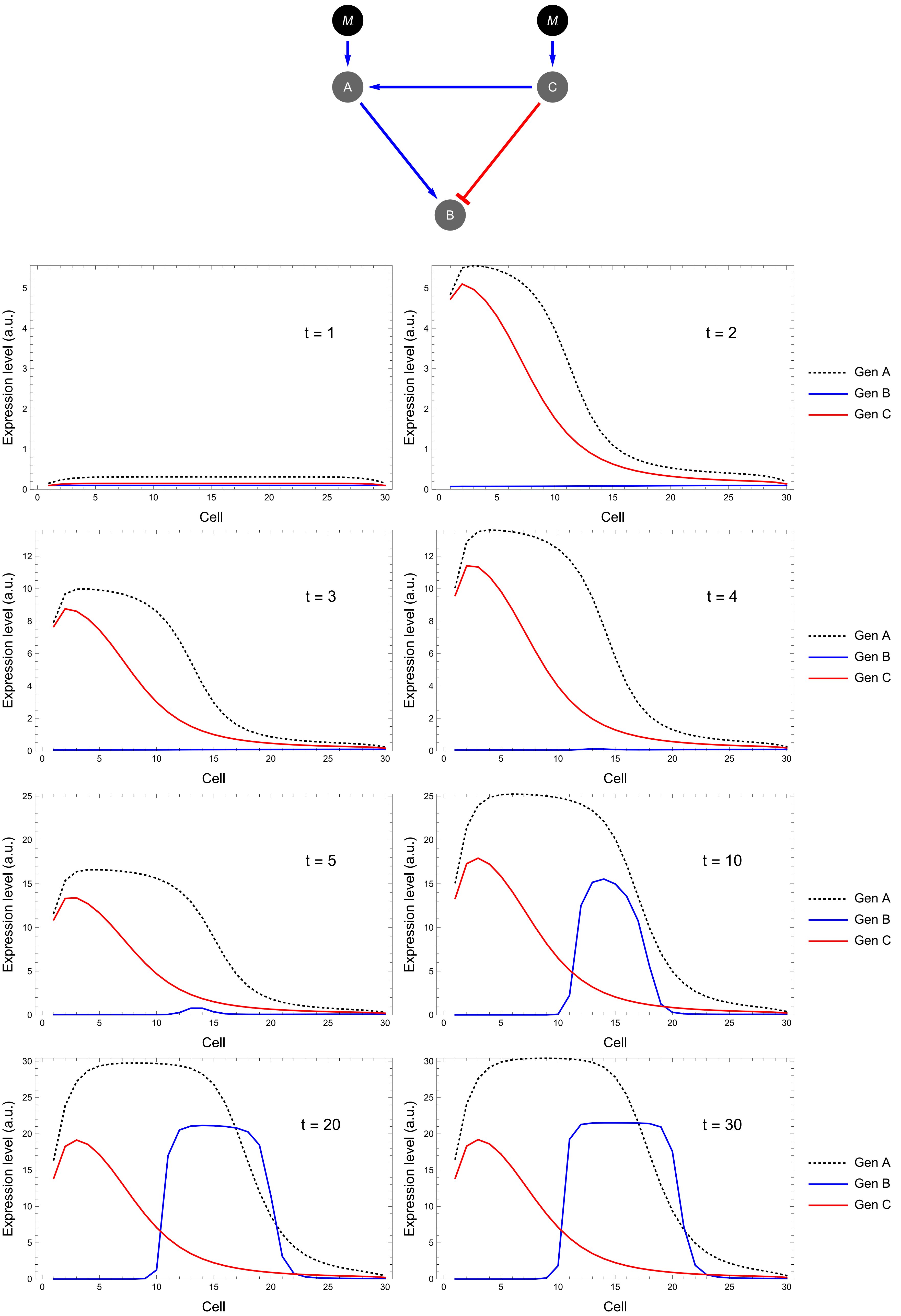

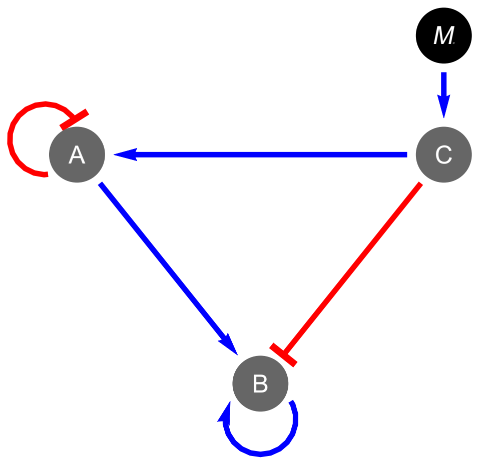

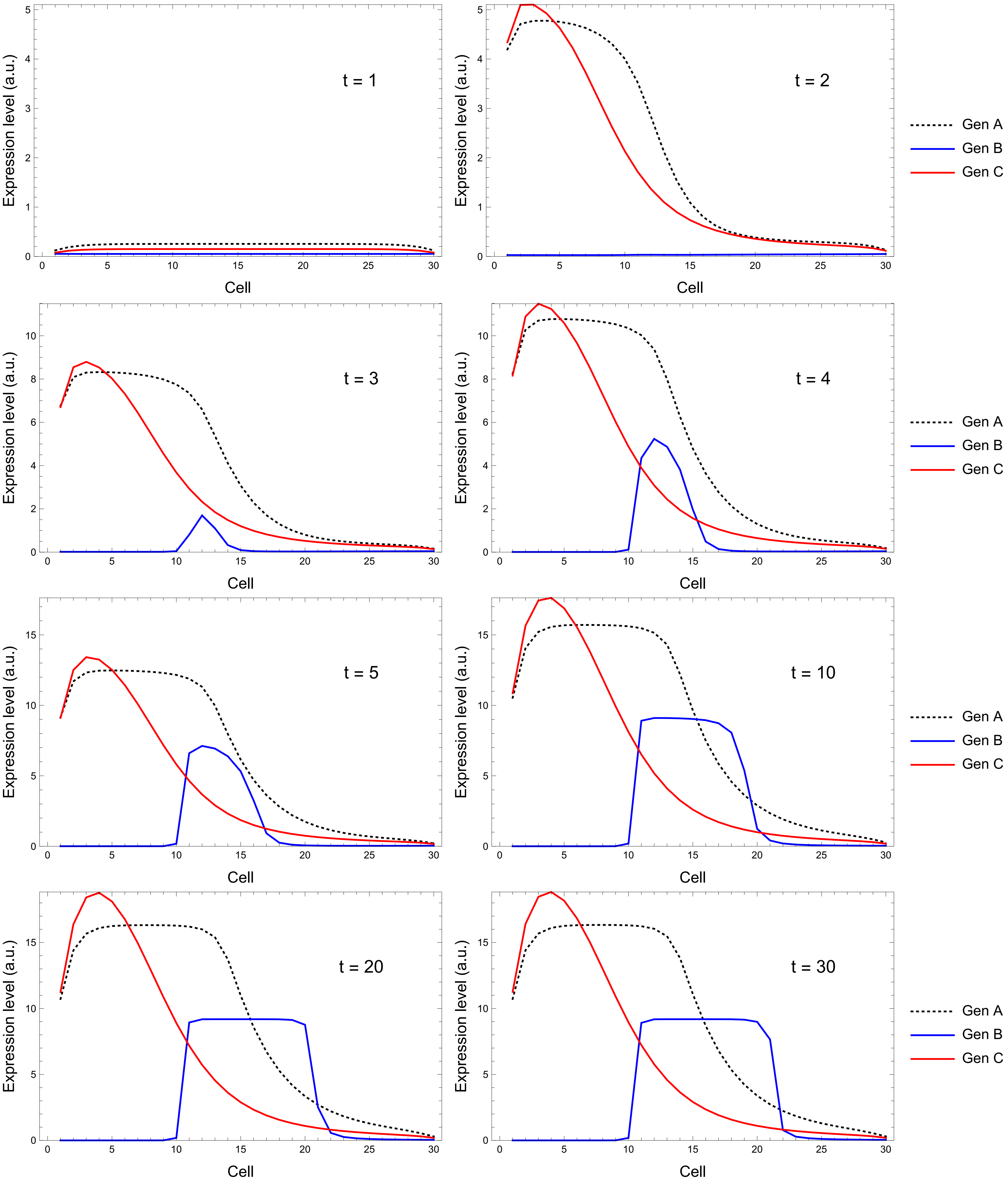

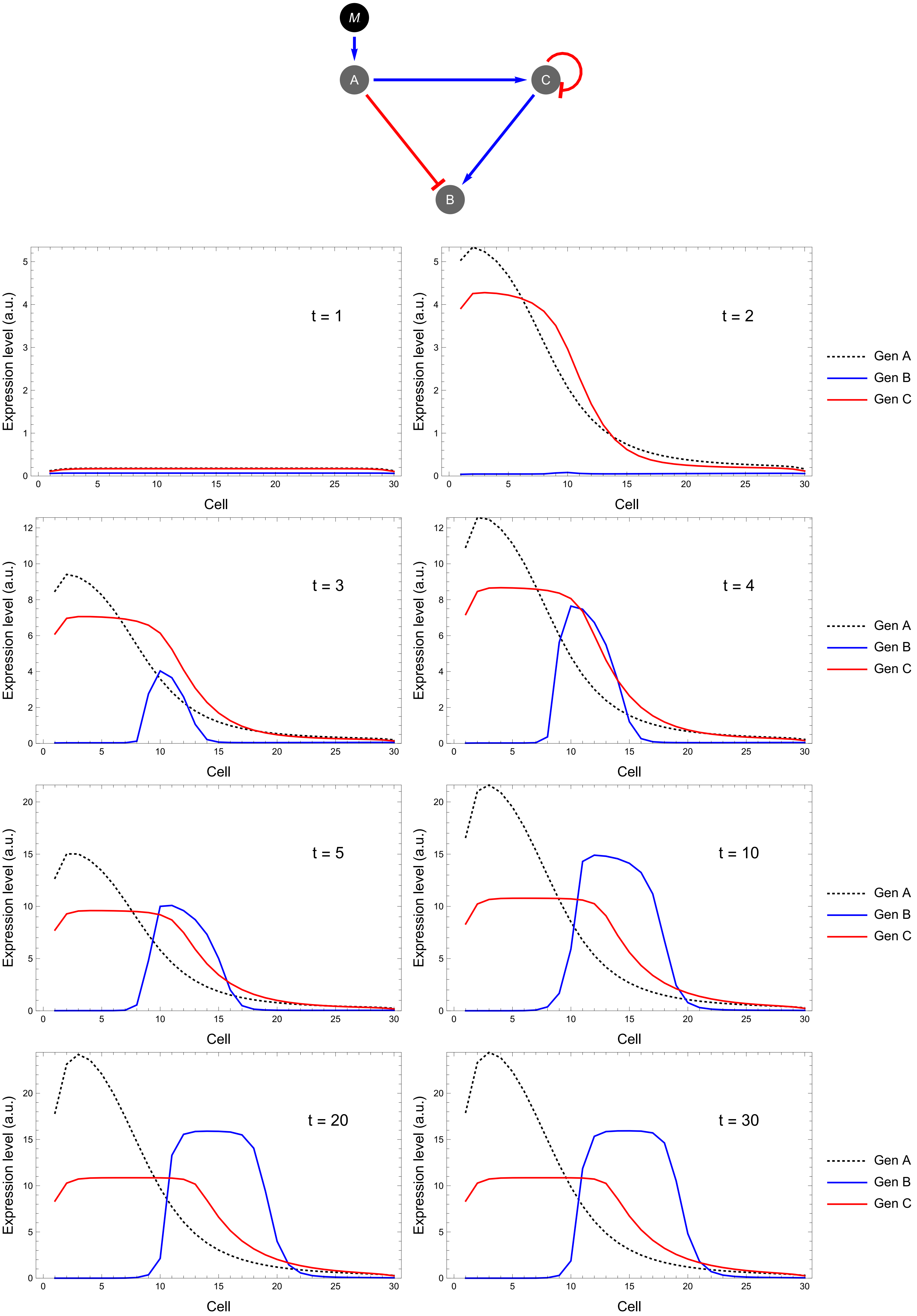

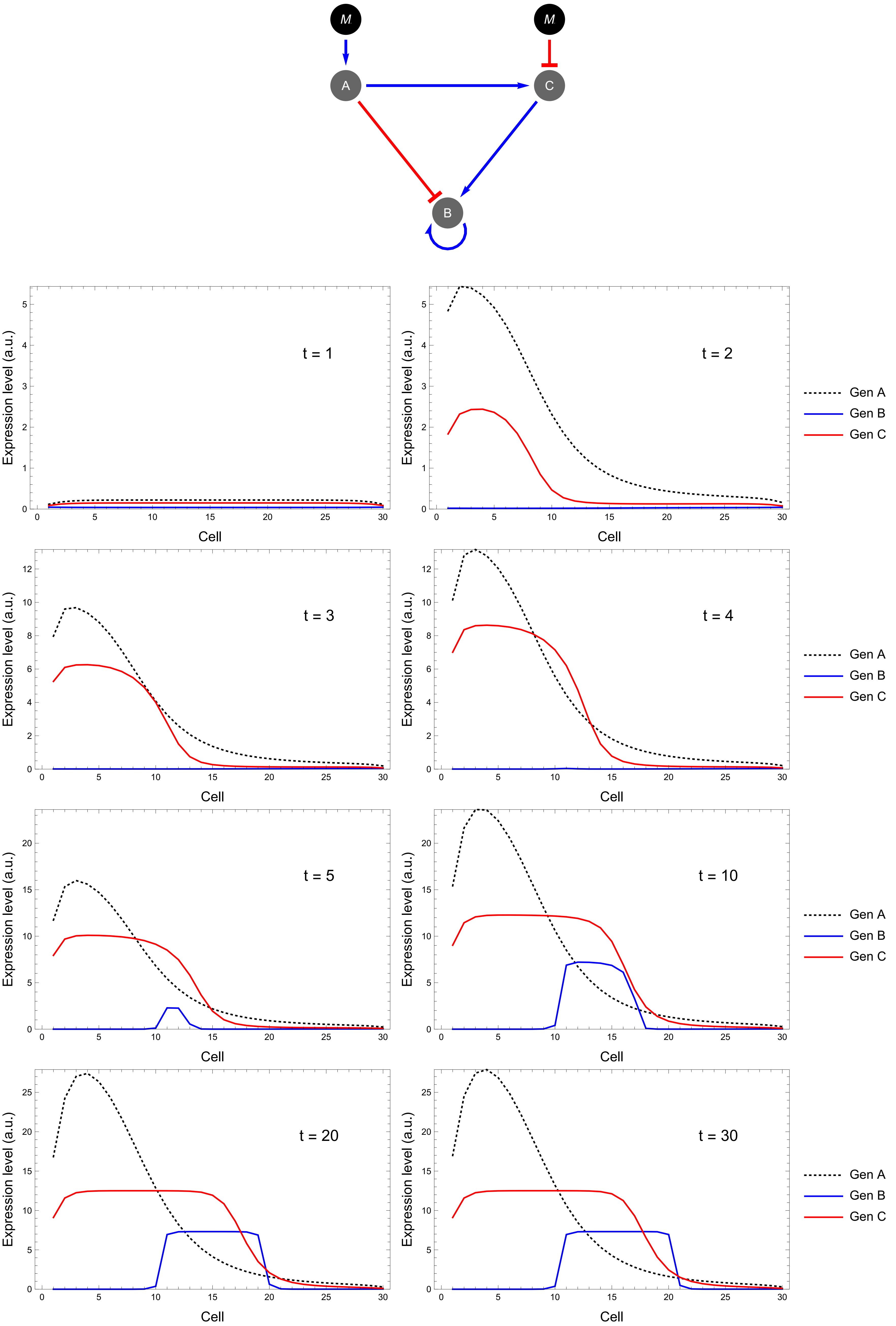

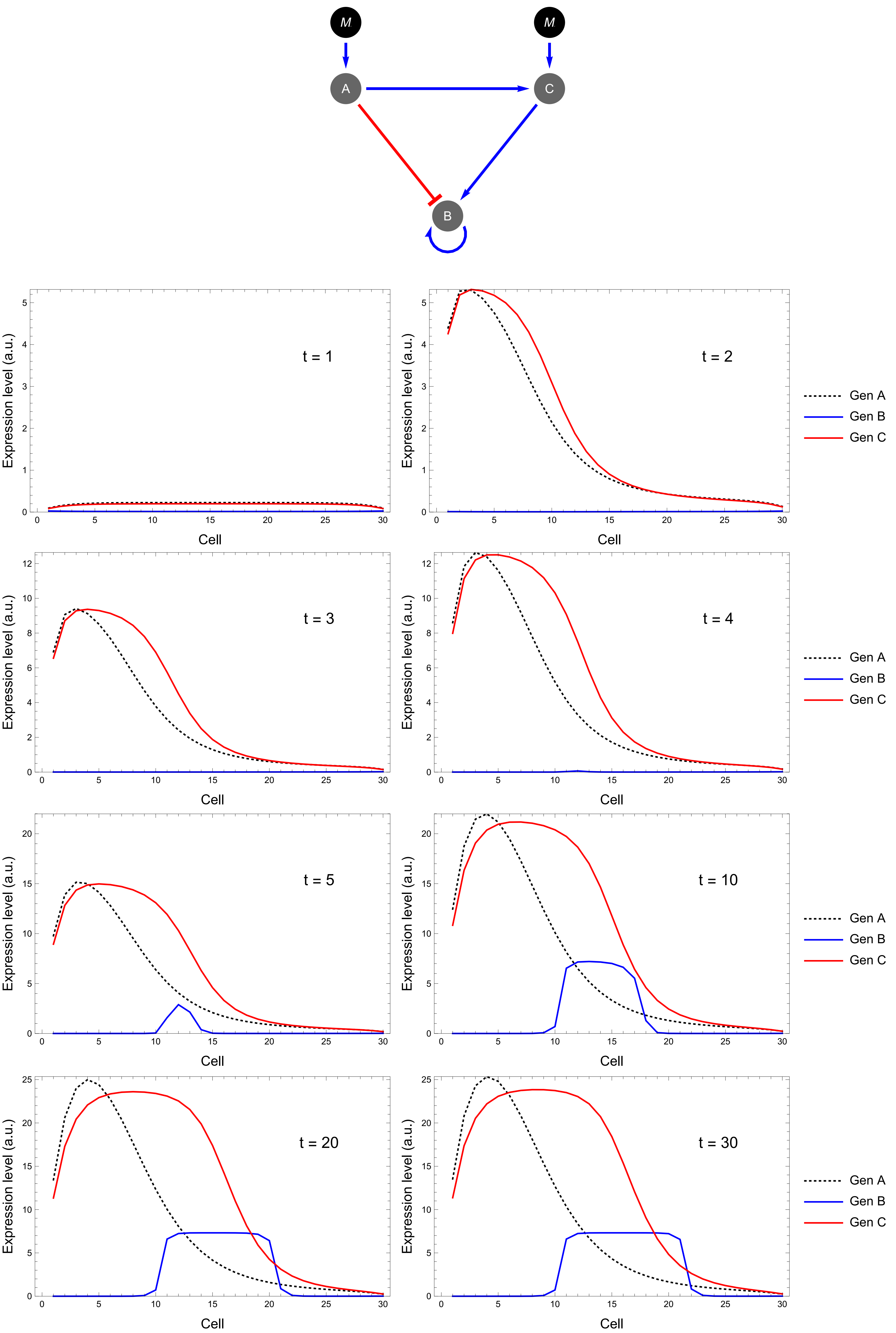

### S5 Fig

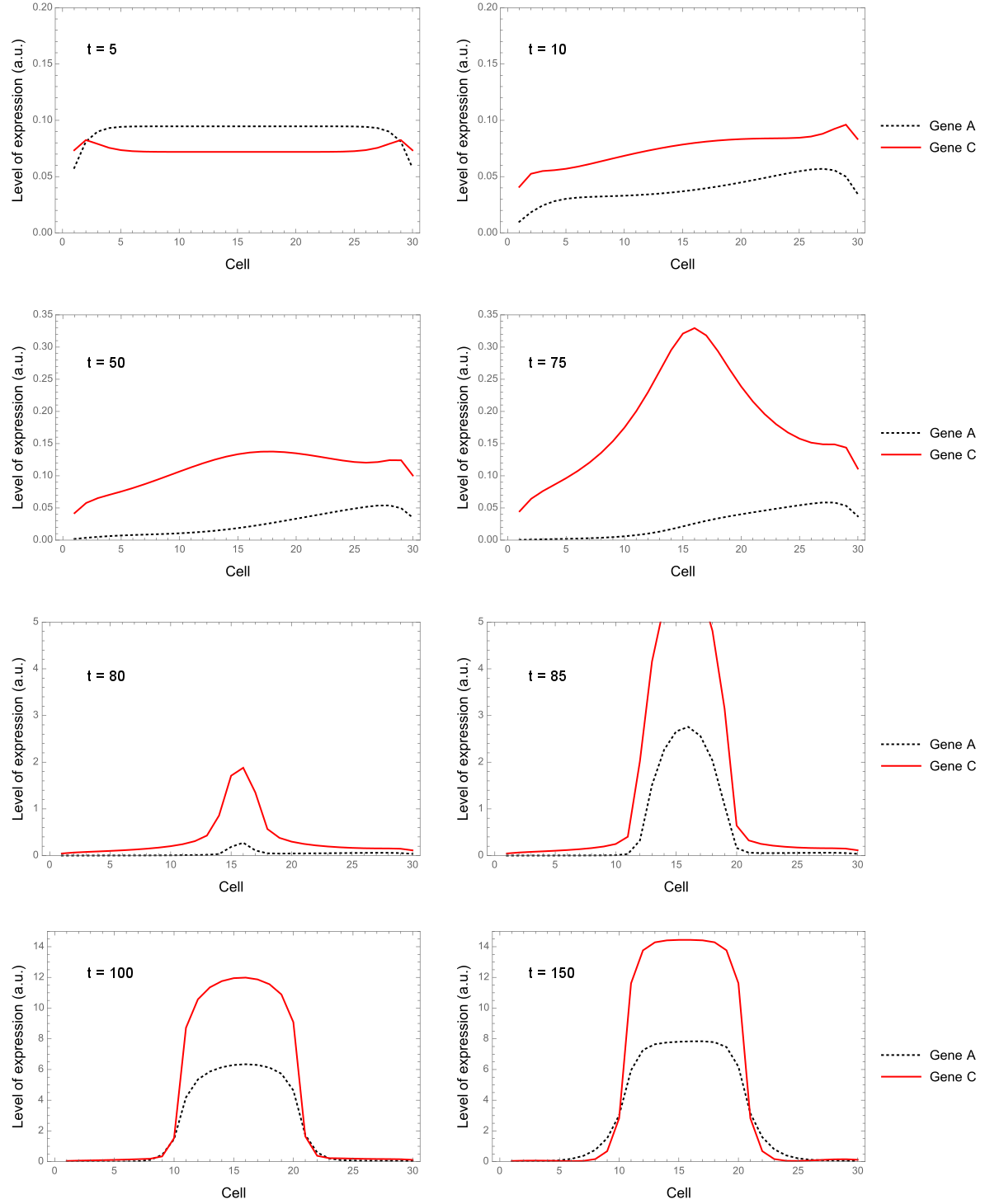

### S6 Fig

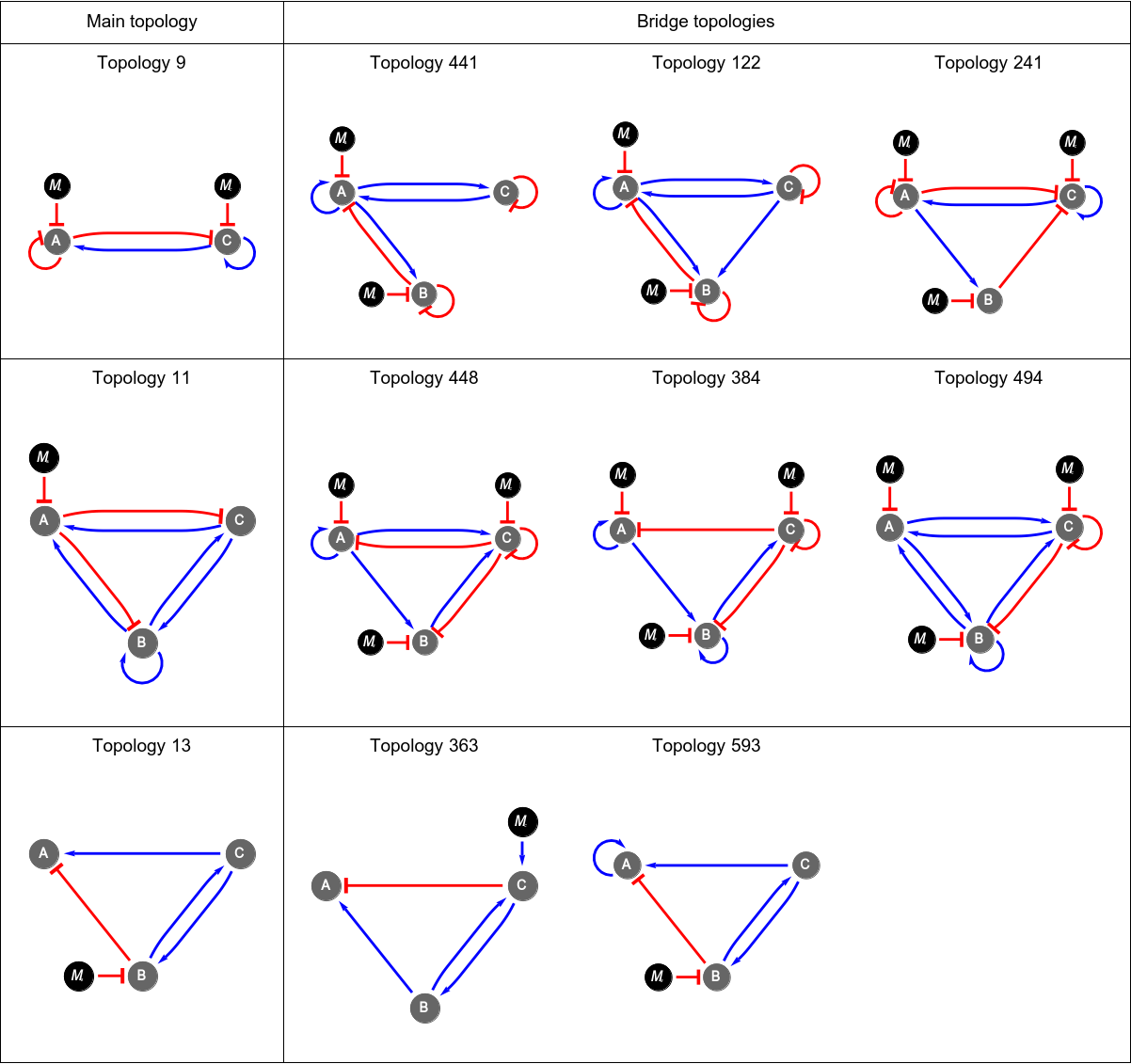

### S7 Fig

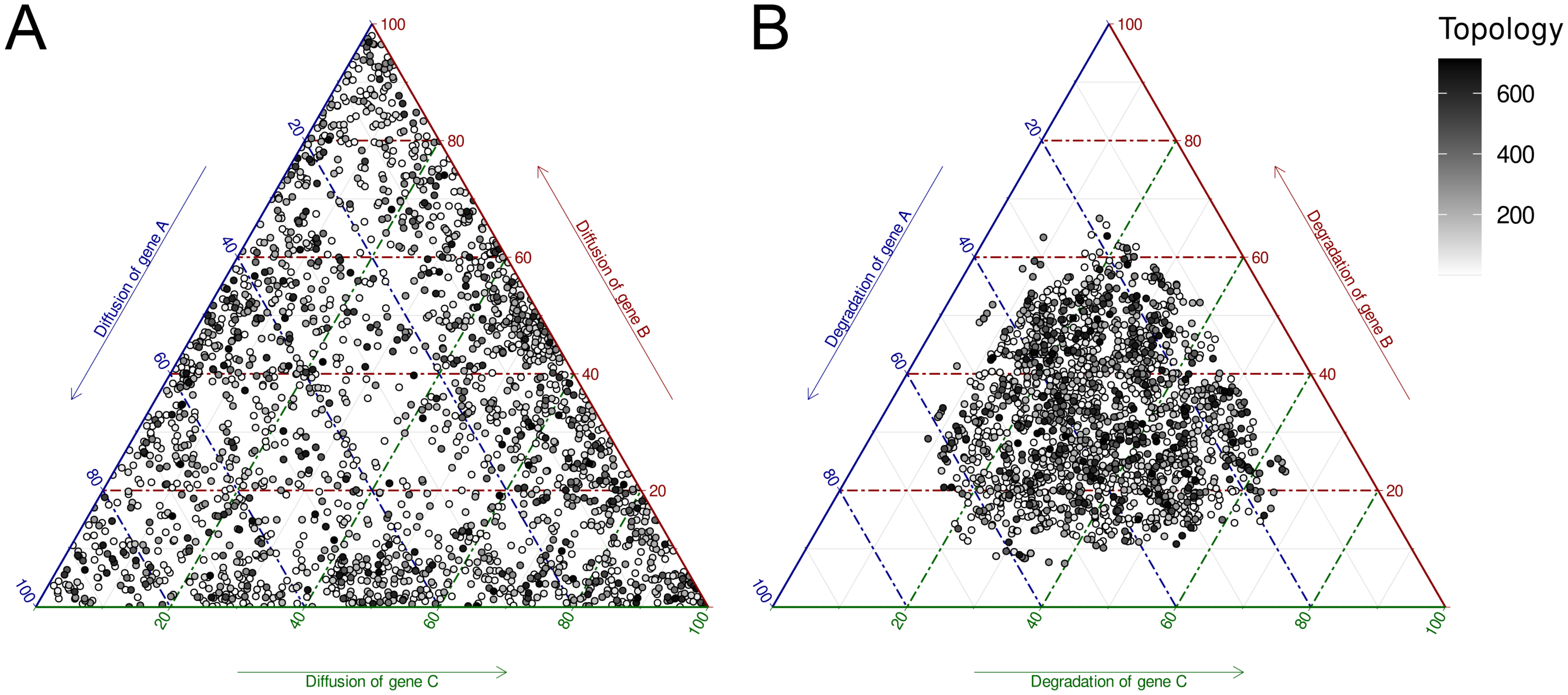
