## Supplementary material for "Elucidating multi-input processing 3-node gene regulatory network topologies capable of generating striped gene expression patterns": S1 Appendix

### Diffusion independent networks

We investigated the ability of the GRNs to produce the optimal pattern without diffusion. For this purpose, the diffusion parameter for each gene in all the GRNs was set to 0 and then the fitness of each GRN was calculated. Those GRNs with $fitness > 0.95$ were considered to be diffusion independent (Fig A1).

Out of 2061 GRNs, 404 were diffusion independent (Fig A2). These GRNs correspond to a total of 159 topologies. The first eigth topologies, which are variations of the Incoherent type 3 Feed-Forward Loop, have at least one diffusion independent GRNs. On the other hand, topology 9 presented no diffusion independent GRNs (Fig A3). This was expected as these networks are composed of a core very similar to the Gierer-Meinhardt model, which is dependent on diffusion of its components.

### Robustness to changes in the signal input

We also investigated if the sampled GRNs were robust to changes in the morphogen input. For each GRN, we generated a set of 100 morphogen gradients by randomly chosing the parameter $A_0$ from a normal distribution with $\mu = 1$ and $\sigma = 0.3$. Then, we calculated the fitness of the GRNs with each one of the different morphogen gradients. As expected, higher fitness values were obtained when $A_0$ was closer to the original value ($A_0 = 1$) (Fig A4).

Interestingly, among the most abundant topologies, topology 9 was found to be very sensible to changes in the signal input (Fig A5), while other topologies were found to be more robust to these changes (Fig A6).

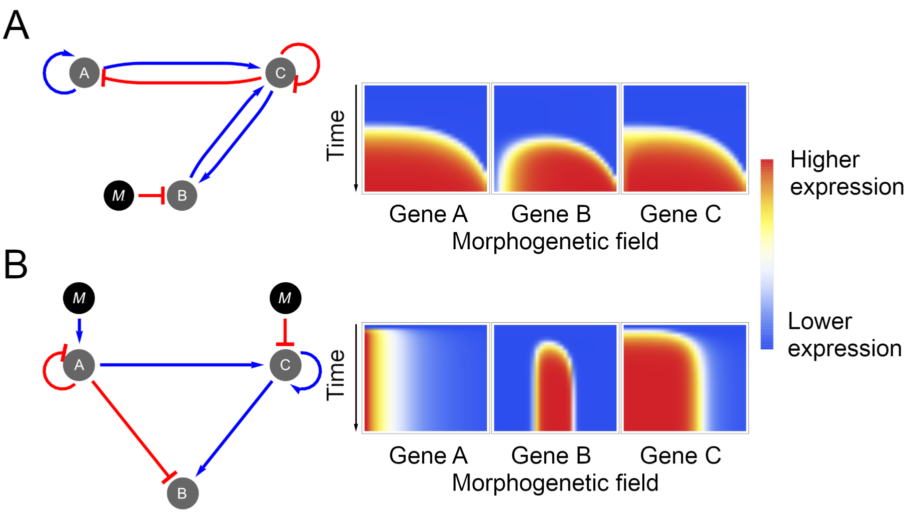


**Fig A1. Diffusion dependent and independent networks.**
(A) Example of a diffusion dependent GRN with its topology (left) and spatio-temporal expression profile when no diffusion is allowed (right). (B) Example of a diffusion independent GRN with its topology (left) and spatio-temporal expression profile when no diffusion is allowed (right).

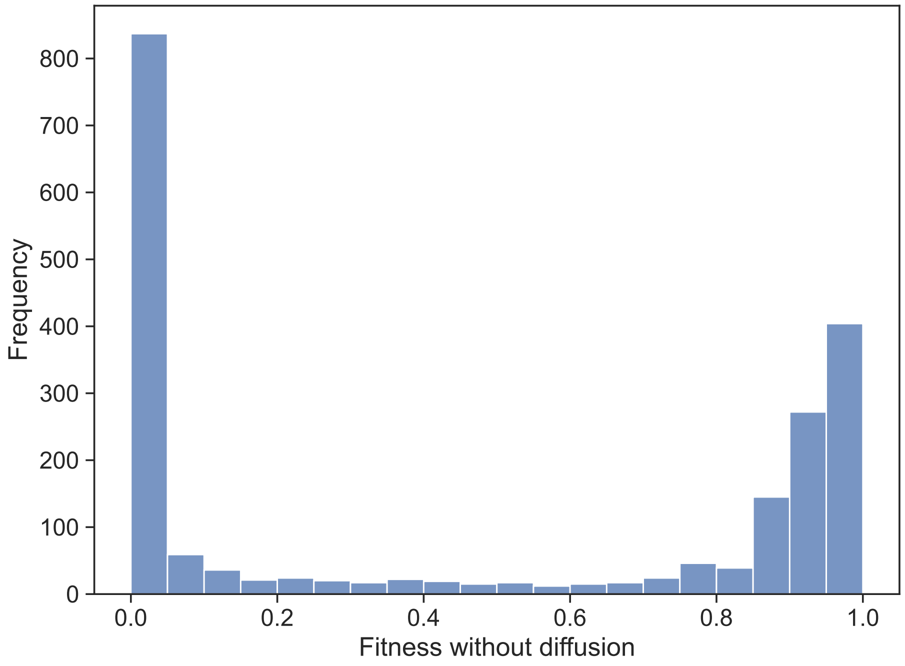


**Fig A2. Histogram of the fitness of GRNs when diffusion is set to 0.**

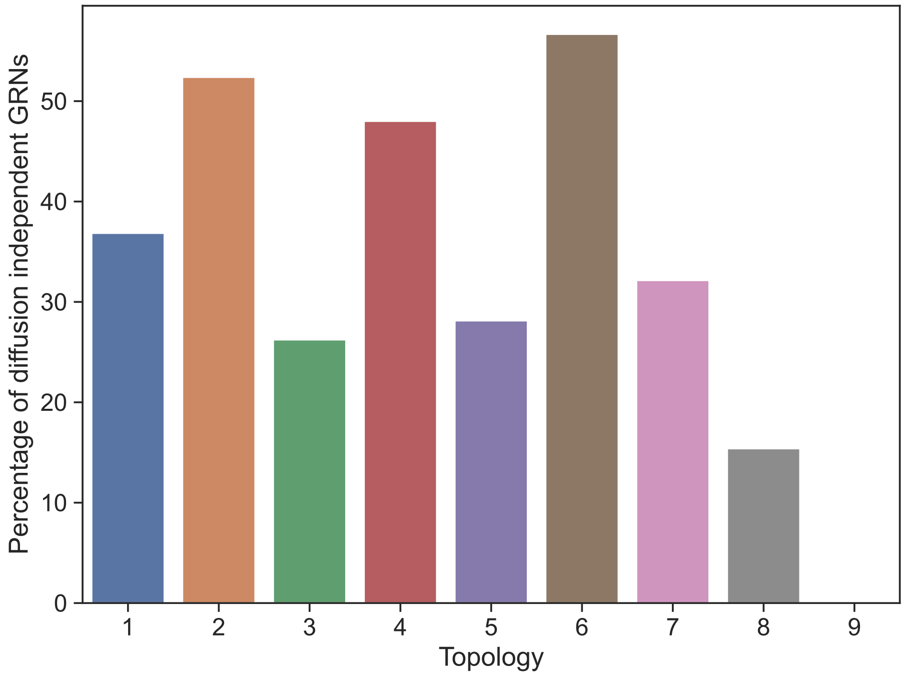


**Fig A3. Percentage of diffusion independent GRNs for the 9 most abundant topologies.**

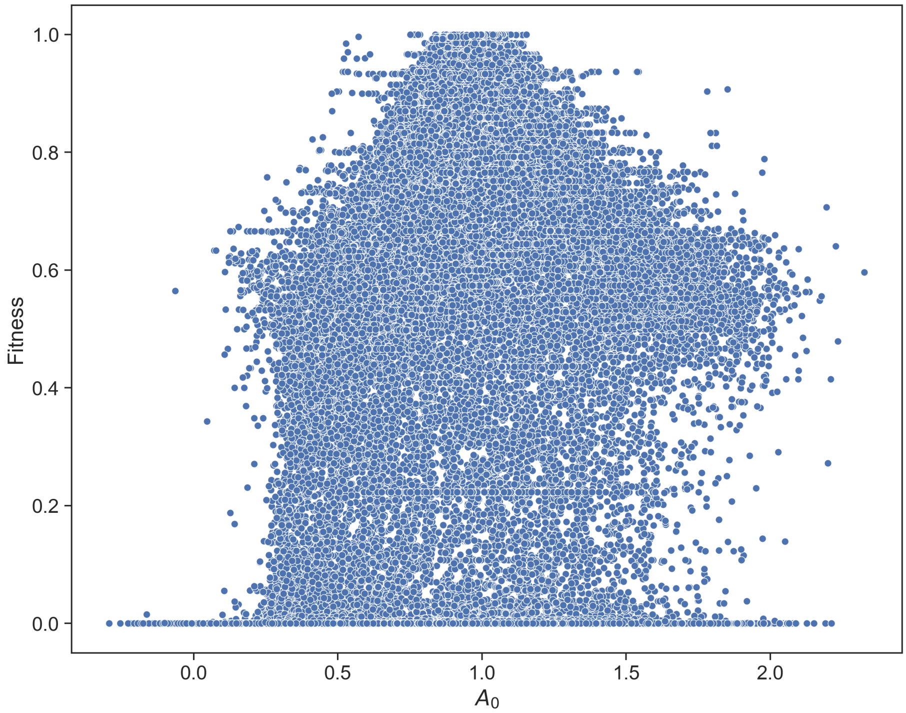


**Fig A4. Robustness of GRNs to random changes in $A_0$.**

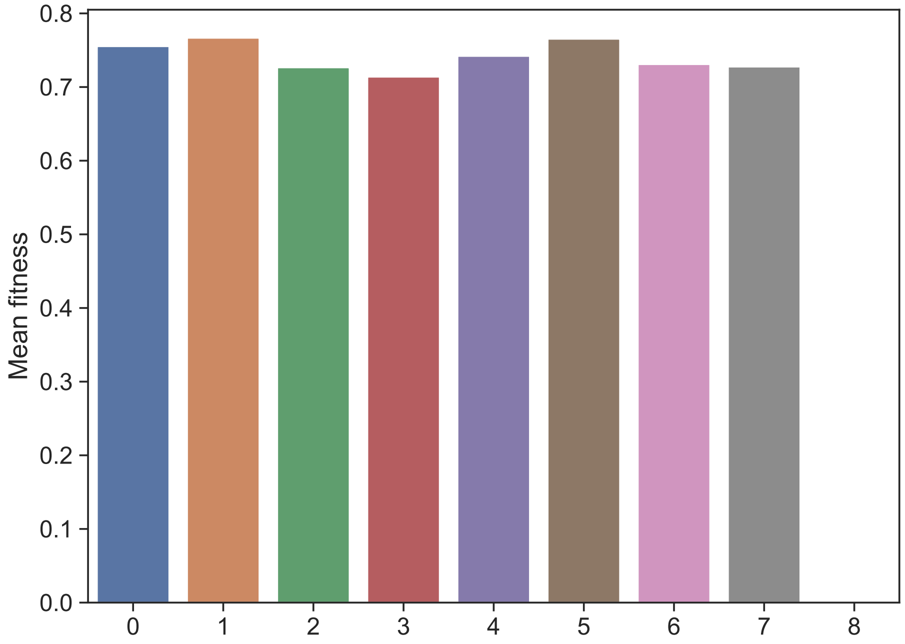


**Fig A5. Mean fitness for the 9 most abundant topologies.**

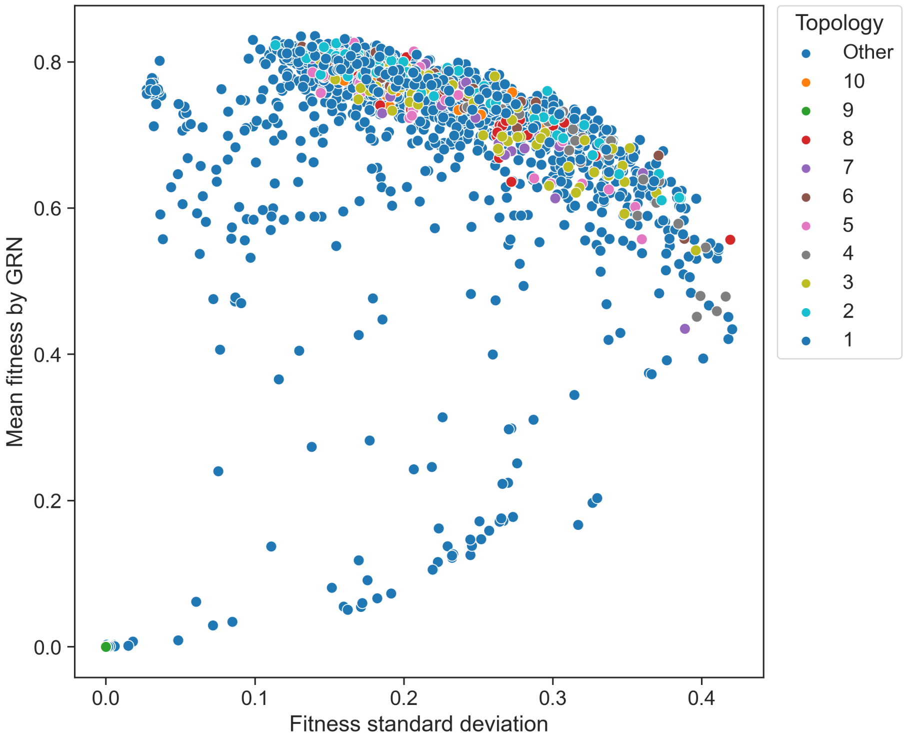


**Fig A6. Mean fitness by GRN vs. Fitness standard deviation.**
